# Overactivation of prefrontal astrocytes impairs cognition through the metabolic pathway of central kynurenines

**DOI:** 10.1101/2024.12.02.626319

**Authors:** Viktor Beilmann, Johanna Furrer, Sina M. Schalbetter, Ulrike Weber-Stadlbauer, Matthias T. Wyss, Aiman S. Saab, Bruno Weber, Urs Meyer, Tina Notter

## Abstract

Astrocyte dysfunctions have long been implicated in psychiatric and cognitive disorders, yet the precise mechanisms underlying this association remain elusive. Here, we show that chemogenetic activation of prefrontal astrocytes in mice impairs short-term memory and sensorimotor gating and attenuates the activation of parvalbumin (PV) interneurons in the prefrontal cortex. These alterations are accompanied by increases in prefrontal levels of kynurenic acid (KYNA), a key metabolite of the kynurenine (KYN) pathway, known to be produced by astrocytes, which serves as an endogenous antagonist of NMDA receptors. Pharmacological inhibition of kynurenine aminotransferase II, the key enzyme mediating the transamination of KYN to KYNA, reinstates the astrocyte-mediated impairments in short-term memory and sensorimotor gating, and normalizes the deficits in prefrontal PV interneuron activation. Our study identifies a mechanistic link between overactivation of prefrontal astrocytes, increased production of KYNA, and cognitive as well as cellular dysfunctions involved in major psychiatric disorders and beyond.

## INTRODUCTION

Astrocytes are macroglial cells of neuroepithelial origin that contribute to a multitude of homeostatic processes in the brain.^1^ Being an integral part of the tripartite synapse^2^, they modulate synaptic activity through multiple actions on neurotransmitter clearance, recycling and release^3^, and act as central metabolic hubs for energy and biosynthesis^4, 5^. Astrocytes sense and respond to neuronal activity through a variety of G-protein coupled receptors (GPCRs).^6^ Activation of astrocytic GPCRs elicits an elevation of their intracellular Ca^2+^ concentration, leading to the release of neuroactive molecules (gliotransmitters), which modulate neuronal activity and synaptic strength.^2, 3, 6^ These molecular processes might offer a means by which astrocytes regulate cognitive functions.^7^

Astrocytic dysfunctions have long been implicated in cognitive^8, 9^ and psychiatric^3, 10-13^ disorders, including (but not limited to) schizophrenia and other psychosis-related disorders. For example, post-mortem evidence suggests that at least a portion of individuals with psychotic disorders display increased cortical expression of cellular markers indicative of astrocyte activation, such as glial fibrillary acidic protein (GFAP)^14-16^, although these findings remain inconclusive^11, 17^. Moreover, several transcriptomic studies have identified increased expression of astrocytic gene sets in prefrontal post-mortem samples obtained from individuals with psychotic disorders^18-22^, which matches recent findings implicating astrocytes to familial risk and clinical symptoms of schizophrenia.^23^ Hence, overactivation of astrocytes may contribute to the pathophysiology of schizophrenia and related psychotic disorders, but the precise mechanisms underlying this association remain elusive.

Such astrocytic dysfunctions might hold significant relevance for cognitive functions, particularly when they manifest within the prefrontal cortex (PFC). The PFC provides executive control of a variety of cognitive processes, including attention, working memory, decision making, and goal-directed behavior, allowing information processing and behavior to vary adaptively depending on current goals.^24^ Functional and structural anomalies of the PFC are also central features of several psychiatric disorders with cognitive impairments, including schizophrenia and other psychosis-related disorders.^25-29^ Prefrontal abnormalities in these disorders include increased glutamate levels^30^, alterations in parvalbumin (PV) interneurons^19, 31, 32^, and reduced *N*-methyl-D-aspartate (NMDA) receptor activation^31, 33^, pointing towards multifaceted alterations in prefrontal glutamatergic and GABAergic neurotransmission. Additionally, prefrontal abnormalities observed in schizophrenia and related disorders involve the presence of increased kynurenic acid (KYNA).^16, 34, 35^ KYNA, a key metabolite of the kynurenine (KYN) pathway of tryptophan degradation^36^, acts as an endogenous antagonist of NMDA receptors and is known to impair various cognitive functions, including short-term memory^36, 37^ and sensorimotor gating^38^. Unlike other glial cells, astrocytes abundantly express the enzyme kynurenine aminotransferase II (KATII)^39^, which mediates the transamination of KYN into KYNA, and therefore, they are a primary source for KYNA production and secretion in the brain parenchyma in health and disease.^36^

Taken together, there is accumulating evidence for astrocytic, metabolic, and neuronal anomalies in prefrontal cortical areas in patients with schizophrenia and other psychosis-related disorders. It remains unknown, however, how these abnormalities relate to one another and how they contribute to specific behavioral and cognitive functions affected in psychosis-related disorders. Establishing a causal link between these cellular and cognitive processes is warranted in order to advance our understanding of the mechanisms by which alterations in astrocytic functions contribute to cognitive impairments in schizophrenia and other psychosis-related disoders^25-29, 16, 34, 35, 40, 41^.

Therefore, the present study examined how selective overactivation of prefrontal astrocytes modulates PFC-related behavioral and cognitive functions in mice. To achieve this goal, we took advantage of the designer receptor exclusively activated by designer drugs (DREADDs) system to selectively manipulate astrocyte activity in the PFC in mice.^42^ Our DREADDs system was based on the astrocytic expression of the muscarinic Gq-protein-coupled receptor (hM3DGq). This construct was selected because activation of the Gq-protein results in the mobilization of cytosolic Ca^2+^ via inositol-3-phosphate (IP3)^43^, a prevalent signaling pathway regulating astrocyte activity^1^. Using this model, we investigated how overactivation of prefrontal astrocytes modulates PFC-related behavioral and cognitive functions, neuronal activity, and KYNA signaling in the PFC. Our study provides evidence that prefrontal astrocytes regulate neuronal activity and cognitive functions by acting on the metabolic pathway of central KYN, which bears relevance for the cognitive impairments implicated in major psychiatric illnesses.

## RESULTS

### Selectivity and effectiveness of hM3DGq-based stimulation of prefrontal astrocytes

The hM3DGq construct encompassing a fluorescent mCherry tag was delivered via bilateral stereotaxic injections of recombinant adeno-associated viruses (AAV9-hgfaABC1D-hM3DGq-mCherry; **Fig. 1a**) into the PFC of 10-week-old mice, which yielded localized expression of the mCherry tag in anterior cingulate, prelimbic and infralimbic subregions of the PFC in adulthood (**Fig. 1b**). Immunohistochemistry and laser scanning confocal microscopy confirmed the selectivity of construct expression in prefrontal astrocytes, as there was a strong co-localization between the mCherry tag and the astrocyte marker S100β (**Fig. 1c**), but not between mCherry and the microglia marker Iba1 (**Supplementary Fig. S1a**). In quantitative analyses, 98% of all hM3DGq-mCherry positive cells were found to be immunoreactive for S100β (**Fig. 1d**). In line with previous findings in striatal astrocytes^44^, we observed a minor leakage of the construct into neurons, as only ∼2% of the hM3Dq-mCherry-positive cells were immunoreactive for NeuN (**Fig. 1d**). Taken together, the selected experimental approach obtained high astrocytic selectivity of construct expression, with negligible leakage of the construct into neuronal cell populations.

**Figure 1.**
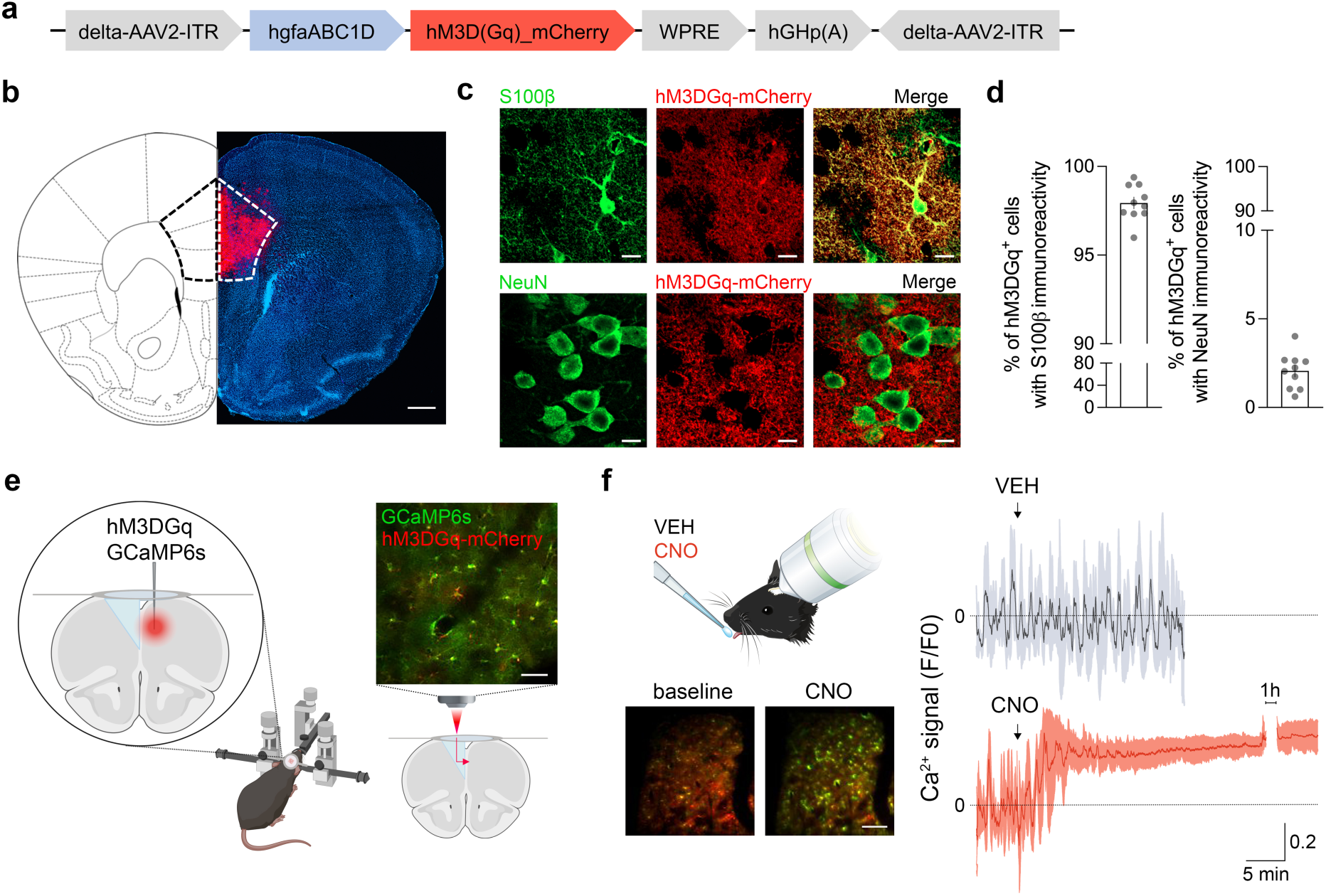
Selectivity and effectiveness of hM3DGq-based activation of prefrontal astrocytes. **(a)** Simplified scheme of the viral vector encompassing hM3DGq under the control of the astrocyte specific promotor hgfaABC1D. **(b)** Bilateral stereotaxic injection of hM3DGq into the PFC resulted in a region-specific expression of the DREADD construct in the target region of interest (medial portion of the PFC; indicated by the dashed line), as shown by the representative tile scan image; hM3DGq expression is shown in red, whereas cell nuclei are depicted in blue (DAPI). Scale bar = 500 μm. **(c)** Representative double-immunofluorescence images confirming cell type selectivity of hM3DGq expression (red) in astrocytes (s100β, green), but not in neurons (NeuN, green). Scale bar = 10 μm. **(d)** Quantification of cellular selectivity of hM3DGq-mCherry construct expression in astrocytes (S100β) and neurons (NeuN). Scatter bar plots represent the percentage of hM3Dq-mCherry-positive cells that were also immunoreactive for the astrocytic marker, S100β (left plot), or the neuronal marker NeuN (right plot) in the PFC. *N* = 10 male mice. **(e)** Surgical procedure for subsequent in vivo two-photon Ca^2+^ imaging of prefrontal astrocytes. rAAVs expressing hM3DGq and GCaMP6s under the astrocyte-specific promoter hgfaABC1D were co-injected into the PFC opposing a right-angled microprism, which was attached to a cranial window implanted into the subdural space within the fissure. The microphotograph shows a representative image of prefrontal astrocytes co-expressing the hM3DGq-mCherry and GCaMP6s construct acquired by in vivo two-photon imaging in awake animals. Scale bar = 25 μm. **(f)** Mode of drug delivery during two-photon Ca^2+^ imaging in awake animals using the MDA approach. The microphotographs show representative images of Ca^2+^ signals (green, GCaMP6s) in hM3DGq-expressing prefrontal astrocytes (red) before (baseline) and after clozapine-N-oxide (CNO, 1 mg/kg) administration. Ca^2+^ responses were indexed by the F/F0 ratio in prefrontal astrocytes after oral vehicle (VEH) or CNO administration via MDA. Black and red traces represent Ca^2+^ responses (means ± SD) to VEH and CNO treatment, respectively. *N* = 4 male mice. Scale bar = 100 μm.

Next, we used in vivo two-photon Ca^2+^ imaging in awake animals to ascertain the effectiveness and temporal dynamics of the hM3DGq-DREADD system in prefrontal astrocytes. To this end, we co-injected the hM3DGq construct with a viral construct (AAV9-hgfaABC1D-GCaMP6s) expressing the Ca^2+^ sensor, GCaMP6s, into one hemisphere of the PFC of adult mice (**Fig. 1e**). To access medial prefrontal cortical areas, a right-angled microprism attached to a cranial window was implanted into the subdural space within the fissure according to established protocols^45^ (**Fig. 1e**, **Supplementary information**), leading to minimal tissue damage and astrogliosis (**Supplementary Fig. S1b**). Indeed, the region of interest for in vivo imaging in awake animals remained intact, while the contralateral regions beneath the microprism were merely compressed (**Supplementary Fig. S1b**). Ca^2+^ imaging in awake mice commenced after the animals were trained in the awake imaging setup. Imaging was performed at baseline and after oral administration of either vehicle (VEH; 0.9% sterile NaCl) or 1 mg/kg clozapine-N-oxide (CNO) via the non-invasive micropipette-guided drug administration (MDA) method^46, 47^ (**Fig 1f**). We found no changes in prefrontal astrocytic Ca^2+^ fluctuations between baseline and after VEH administration (**Fig 1f**). Compared to VEH treatment, however, administration of CNO markedly increased intracellular Ca^2+^ levels, starting approximately 5 min after CNO treatment and remaining elevated for at least 90 minutes (**Fig 1f**; **Supplementary Video 1_VEH; Supplementary Video 2_CNO**). Additional imaging sessions were conducted 2 days after CNO administration to ensure astrocyte recovery and to assess the response toward a second CNO challenge. Astrocyte Ca^2+^ fluctuations were indistinguishable from the first imaging session in terms of their baseline activity and response toward a second CNO treatment (**Supplementary Fig. S1c**). Together, these findings confirm the effectiveness of hM3DGq-based stimulation of prefrontal astrocytes in vivo.

### Chemogenetic activation of prefrontal astrocytes impairs short-term memory and pre-attentive filtering

We next explored whether overactivation of prefrontal astrocytes affects behavioral and cognitive functions. For this purpose, 10-week-old male mice received a bilateral stereotaxic injection of hM3DGq in the PFC and were subjected to behavioral and cognitive testing at 12 weeks of age after acute CNO (1 mg/kg) or VEH administration (**Fig. 2a**). First, we assessed whether activation of prefrontal astrocytes alters innate anxiety-like behavior or basal locomotor activity. For this purpose, the animals were tested in the light-dark box (**Fig. 2b**) and open field tests (**Fig. 2c**). There were no group differences in either test (**Fig. 2b,c**), indicating that overactivation of prefrontal astrocytes did not affect innate anxiety or basal locomotor activity.

**Figure 2.**
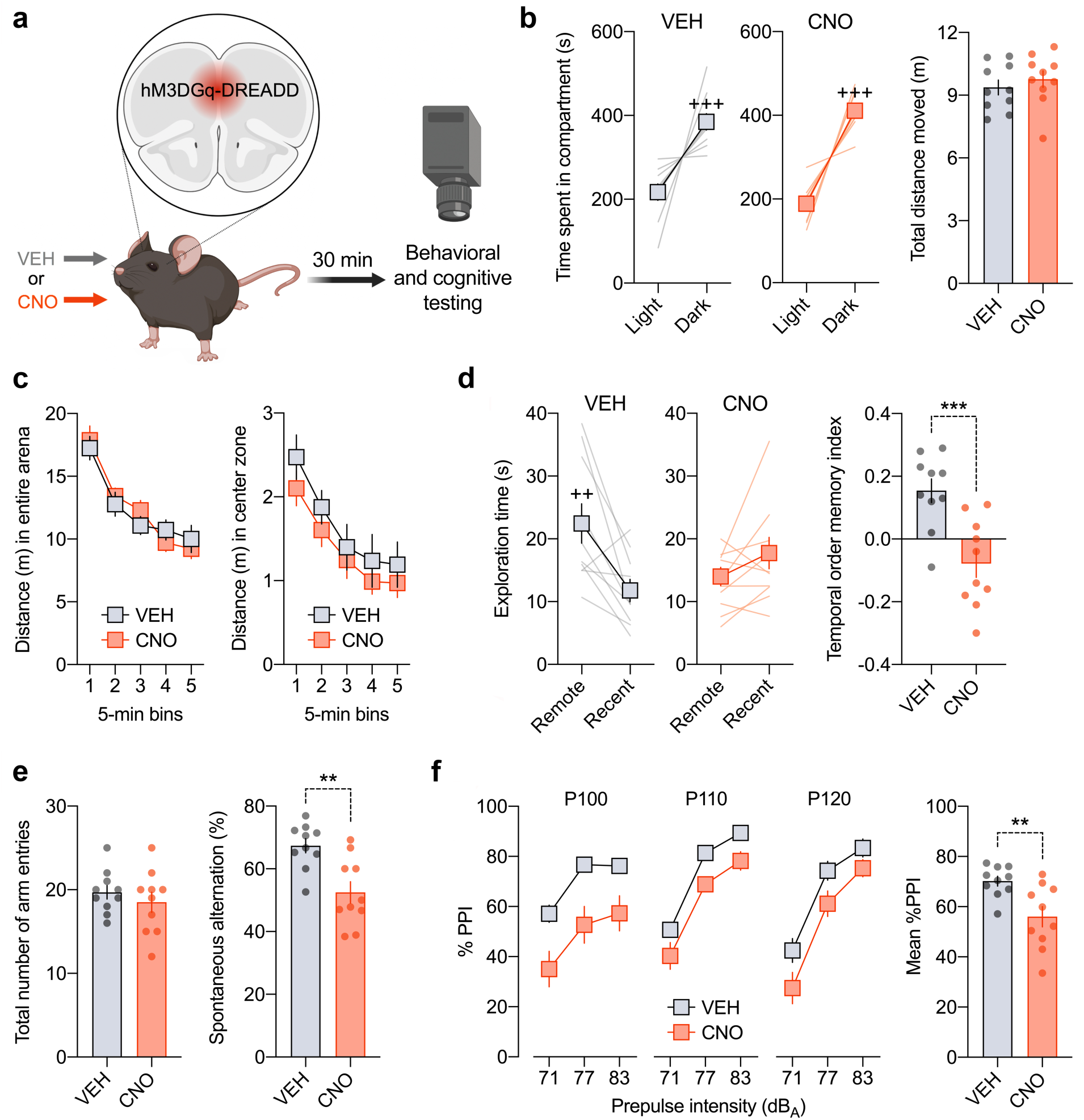
Overactivation of prefrontal astrocytes impairs short-term memory and pre-attentive filtering in male mice. **(a)** Male mice expressing hM3DGq in the prefrontal cortex were treated with vehicle (VEH) or clozapine-N-oxide (CNO, 1 mg/kg) and were then subjected to behavioral and cognitive testing 30 min after treatment. **(b)** Time spent in the light and dark compartments (line plots) and total distance moved (box plot) in the light-dark box test of innate anxiety-like behavior. ^+++^*p* < 0.001, reflecting the significant main effect of compartment revealed by repeated-measure ANOVA (*F*_(1,18)_ = 67.5). **(c)** Distance moved as a function of 5-min bins in the entire arena (left) and center zone (right) during the open field test of exploratory activity. **(d)** Absolute exploration times of the temporally remote and recent objects (line plots) and temporal order memory index (bar plot) in the temporal order memory test for objects. ^++^*p* < 0.01, reflecting the significant main effect of object in VEH-treated mice revealed by repeated-measure ANOVA (*F*_(1,9)_ = 12.3); \*\*\**p* < 0.001, based on two-tailed t-test (*t*_(18)_ = 4.01). **(e)** Total number of arm entries (left) and percent spontaneous alternation (right) in the Y-maze test of working memory. \*\**p* < 0.01, based on two-tailed t-test (*t*_(18)_ = 3.61). **(f)** Prepulse inhibition (PPI) test of pre-attentive filtering. The line plots show % PPI as a function of prepulse intensity (71, 77 and 83 dB_A_) for each of the three pulse conditions (P100, P110 and P120, which correspond to pulse intensities of 100, 110 and 120 dB_A_). The bar plot depicts the mean % PPI across all prepulse and pulse intensities. \*\**p* < 0.01, reflecting the significant main effect of treatment revealed by repeated-measure ANOVA (*F*_(1,18)_ = 9.98). All data are means ± SEM with individual values overlaid; *N* = 10 male mice per group and test.

We then examined whether overactivation of prefrontal astrocytes alters cognitive functions. For this purpose, we subjected the mice to the temporal order memory for objects test. In this task, which is dependent on the integrity of the PFC^48^, mice were first allowed to freely explore a first pair of identical objects (sample phase 1) and then a novel pair of identical objects (sample phase 2). In the subsequent test phase, mice with intact PFC functions are expected to be able to discriminate between the temporally remote and the temporally recent object, spending more time exploring the object from the first phase. While VEH-treated hM3DGq mice showed a clear preference toward the temporally remote object, CNO-treated hM3DGq mice failed to discriminate between the two objects (**Fig. 2d**). Notably, the amount of time spent exploring the objects in sample phases 1 and 2 was comparable between the groups (phase 1: 71.4 ± 6.5 s (mean ± SEM) for VEH- and 81.6 ± 6.6 s for CNO-treated mice; phase 2: 63.1 ± 5.6 s for VEH- and 70.8 ± 9.0 s for CNO-treated mice). Thus, the disruption of temporal order memory after overactivation of prefrontal astrocytes represented a selective deficit in the capacity to discriminate the relative recency of stimuli^48^. Consistent with these findings, overactivation of prefrontal astrocytes also impaired working memory in the form of spontaneous alternation in the Y-maze, in which performance is dependent on PFC^49^. We found that CNO-induced activation of prefrontal astrocytes in hM3DGq mice led to a significant reduction in % alternation scores without affecting the total number of arm entries (**Fig. 2e**), demonstrating selective effects on working memory^49^.

Next, we investigated whether overactivation of prefrontal astrocytes affects pre-attentive filtering in the form of prepulse inhibition (PPI) of the acoustic startle reflex, which refers to the attenuation of the reflexive startle response to an intense acoustic pulse stimulus when its presentation is shortly preceded by a weak prepulse stimulus^50^. PPI correlates with certain cognitive functions^51^ and is deficient in various neuropsychiatric disorders, including schizophrenia and related disorders^50^. As shown in **Fig. 2f**, prefrontal astrocyte activation led to a significant overall reduction in % PPI, which was evident across all pulse intensities investigated. There were no group differences in terms of prepulse-induced reactivity (mean ± SEM in VEH group: 32.2 ± 4.9 [arbitrary units, AU]; mean ± SEM in CNO group: 30.6 ± 4.2 [AU]) or startle reactivity (mean ± SEM in VEH group: 185.8 ± 29.3 [AU]; mean ± SEM in CNO group: 199.2 ± 34.7 [AU]), indicating that the reduction in % PPI emerging after overactivation of prefrontal astrocytes reflects genuine deficits in pre-attentive filtering.

To exclude non-selective effects of CNO^52^, we repeated the same behavioral and cognitive testing battery in male mice that received stereotaxic injections of a control AAV9 (hereinafter referred to as ConV), which only expressed a fluorescent EGFP tag under the same GFAP promoter (**Supplementary Fig. S2**). We found that VEH- and CNO-treated ConV mice did not differ in any of the behavioral or cognitive measures of interest, suggesting that the cognitive changes emerging in CNO-treated hM3DGq mice (**Fig. 2**) represent genuine effects induced by overactivation of prefrontal astrocytes.

Furthermore, we examined the same behavioral and cognitive functions in female mice expressing hM3DGq in prefrontal astrocytes (**Supplementary Fig. S3**) in order to identify possible sex-dependent effects. Consistent with males, CNO-treated female mice expressing hM3DGq did not differ in terms of innate anxiety-like behavior (**Supplementary Fig. S3b**) or basal locomotor activity (**Supplementary Fig. S3c**). Moreover, consistent with the effects in males, overactivation of prefrontal astrocytes in female mice impaired the capacity to discriminate the relative recency of stimuli in the temporal order memory test for objects (**Supplementary Fig. S3d**) and working memory in the Y-maze spontaneous alternation test (**Supplementary Fig. S3e**). Although somewhat less extensive than males, female mice expressing hM3DGq also displayed significant deficits in % PPI of the acoustic startle reflex test after CNO treatment (**Supplementary Fig. S3f**), specifically under test conditions in which 120 dB_A_ stimuli served as pulses. Together, these findings demonstrate that overactivation of prefrontal astrocytes similarly disrupts short-term memory and pre-attentive filtering in male and female mice without affecting indices of anxiety-like behavior or locomotor activity.

### Overactivation of prefrontal astrocytes alters neuronal activation in the PFC

To explore the mechanisms by which prefrontal astrocytes modulate cognitive functions, we first examined whether overactivation of prefrontal astrocytes affects neuronal activation patterns in the PFC of mice undergoing temporal order memory testing. For this purpose, hM3DGq expressing male mice treated with either VEH or CNO were sacrificed 2 hrs after completion of the temporal order memory test, after which brains were collected for subsequent immunohistochemical analyses (**Fig. 3a**). As before^53^, immunoreactivity for the immediate early gene c-Fos was hereby taken as a proxy for neuronal activation. Prior to these analyses, we confirmed the deficit in temporal order memory in CNO-treated hM3DGq mice relative to VEH-treated controls (**Fig. 3b**). In CNO-treated hM3DGq mice with reduced temporal order memory, we identified an overall increase in the number of c-Fos positive neurons in the PFC (**Fig 3c**). We next quantified c-Fos expression specifically in PV interneurons (**Fig. 3d**), given their known functions in supporting PFC-dependent working memory^54^, as well as their implication in psychiatric disorders with cognitive impairments^19, 31, 32^. Strikingly, we found a significant reduction in the percentage of c-Fos positive PV interneurons in CNO-treated hM3DGq mice compared to controls (**Fig. 3e**). This reduction emerged in the absence of concomitant effects on the total number of PV interneurons (**Fig. 3e**), demonstrating that overactivation of prefrontal astrocytes induces selective effects on the activation of PV interneurons.

**Figure 3.**
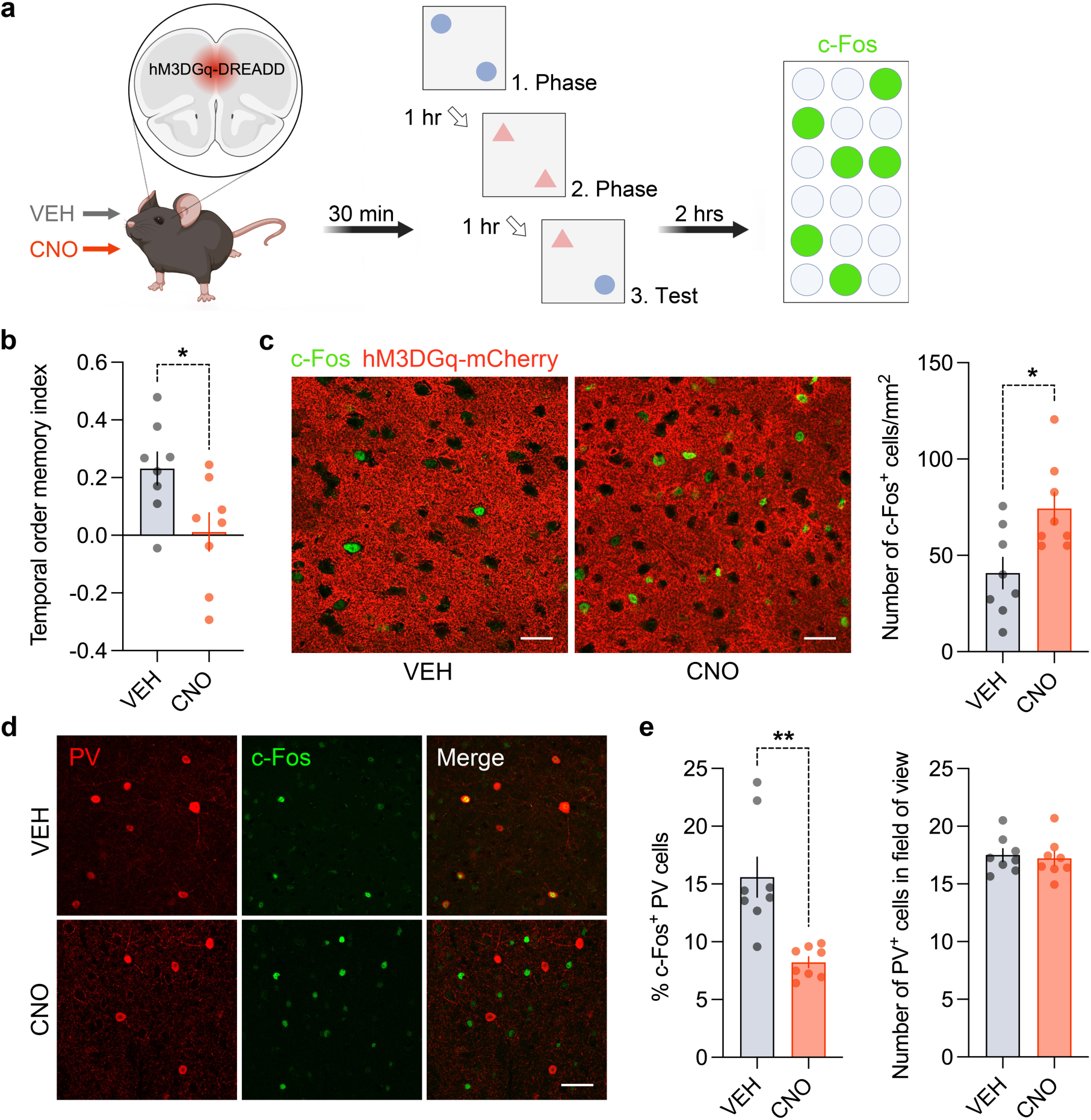
Overactivation of prefrontal astrocytes alters neuronal activation patterns in the PFC. **(a)** Male mice expressing hM3DGq in prefrontal astrocytes were treated with vehicle (VEH) or clozapine-N-oxide (CNO, 1 mg/kg) and were then subjected to the temporal order memory for objects test 30 min after treatment. Two hours after completion of the test, the animals were sacrificed, and brains were collected for subsequent c-Fos expression analysis using immunohistochemistry. **(b)** Temporal order memory index in the temporal order memory test for objects. \**p* < 0.05, based on two-tailed t-test (*t*_(14)_ = 2.52). **(c)** The photomicrographs depict representative immunofluorescent images of mCherry (red) and c-Fos (green) in the PFC of VEH-treated and CNO-treated hM3DGq mice. Scale bars = 25 μm. Quantification of c-Fos positive cells per mm^2^ in the PFC of VEH-treated vs CNO-treated animals. \**p* < 0.05, based on two-tailed t-test (*t*_(14)_ = 2.904). **(d)** The photomicrographs depict representative immunofluorescent stainings against parvalbumin (PV, red) and c-Fos (green) in the PFC of VEH-treated and CNO-treated hM3DGq-expressing mice. Co-localization between c-Fos and PV appears yellow (merged image). Scale bar = 25 μm. The bar plots depict % of c-Fos positive PV interneurons and the total number of PV interneurons in the field of view after VEH or CNO treatment. \*\**p* < 0.01, based on two-tailed t-test (*t*_(14)_ = 4.13). All data are means ± SEM with individual values overlaid; *N* = 8 male mice per group and test.

### The central kynurenine pathway mediates the cognitive deficits induced by overactivation of prefrontal astrocytes

In a next step, we aimed to identify the molecular mechanism by which overactivation of prefrontal astrocytes affects neuronal activation and cognitive functions. We hereby focused on the possible involvement of the metabolic pathway of central KYN in general, and the role of KYNA in particular (**Fig. 4a**). Our rationale of focusing on the KYN pathway and KYNA was manifold. First, KYNA acts as an endogenous antagonist at the NMDA receptor and is known to impair short-term memory^36, 37^ and PPI of the acoustic startle reflex^38^, akin to what we observed after overactivation of prefrontal astrocytes. Second, subanesthetic doses of synthetic NMDAR antagonists, such as ketamine, impair working memory and PPI^55^ by reducing the activation of PV interneurons in the PFC^56^, similar to what we observed here. Third, astrocytes abundantly express KATII^39^, which mediates the conversion of KYN into KYNA (**Fig. 4a**), and therefore, they are a primary source for KYNA production and secretion in the brain parenchyma^36^. Fourth, because the enzymatic activity of KAT enzymes is enhanced by Ca^2+ 57^, we hypothesized that chemogenetically induced elevation of astrocytic Ca^2+^ levels would increase prefrontal KYNA levels. In agreement with this hypothesis, liquid chromatography-nanoelectrospray ionization tandem mass spectrometry revealed increased KYNA levels in PFC tissue of CNO-treated hM3DGq mice relative to VEH-treated controls (**Fig. 4b**). While KYN levels were not significantly different between groups, the KYNA/KYN ratio in the PFC increased by 2-folds after chemogenetic activation of prefrontal astrocytes (**Fig. 4b**). Other key metabolites of the KYN pathway, including tryptophan, 3-hydroxykynurenine and quinolinic acid, were not affected following astrocyte activation (**Supplementary Fig. S4**). These findings demonstrate that stimulation of prefrontal astrocytes leads to a selective shift in the metabolic pathway of central KYN towards increased KYNA production.

**Figure 4.**
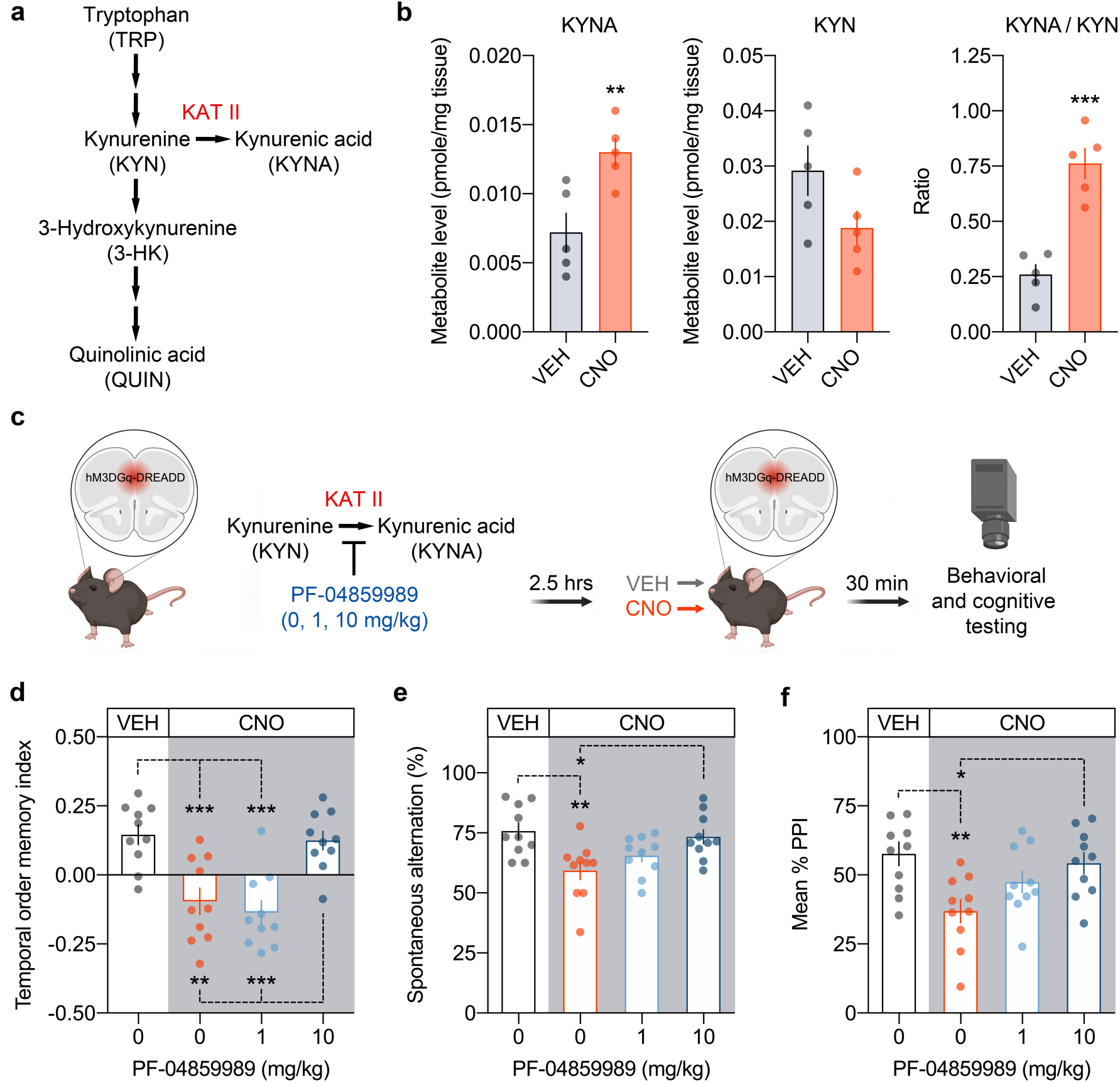
Involvement of the metabolic pathway of central KYN after overactivation of prefrontal astrocytes. **(a)** Simplified scheme of the kynurenine pathway, which generates kynurenic acid (KYNA) through transamination of kynurenine (KYN) by kynurenine aminotransferase II (KATII). **(b)** Levels of KYNA and KYN, as well as the KYNA/KYN ratio, in the PFC of hM3DGq male mice after vehicle (VEH) or clozapine-N-oxide (CNO, 1 mg/kg) treatment. Each data point represents the pooled samples of two mice, with 5 replications for each group and measurement. (c) Schematic representation of the pharmacological rescue study. Male mice expressing hM3DGq in prefrontal astrocytes were pretreated with the KAT II inhibitor PF-04859989 (0, 1, or 10 mg/kg, i.p.) 2.5 hours before receiving CNO or VEH, and 3 hours before they were subjected to behavioral and cognitive testing. (d) Temporal order memory index for CNO-activated hM3DGq mice receiving 0, 1, or 10 mg/kg PF-04859989 relative to VEH-treated hM3DGq mice receiving vehicle (0 mg/kg) only. \*\**p* < 0.01 and \*\*\**p* < 0.001, based on Tukey’s post-hoc test following one-way ANOVA (*F*_(3,36)_ = 12.80, *p* < 0.001). **(e)** Percent spontaneous alternation in the Y-maze test of working memory for CNO-activated hM3DGq mice receiving 0, 1, or 10 mg/kg PF-04859989 relative to VEH-treated hM3DGq mice receiving vehicle (0 mg/kg) only. \**p* < 0.05 and \*\**p* < 0.01, based on Tukey’s post-hoc test following one-way ANOVA (*F*_(3,36)_ = 5.47, *p* < 0.01). **(f)** Mean percent prepulse inhibition (mean % PPI) in CNO-activated hM3DGq mice receiving 0, 1, or 10 mg/kg PF-04859989 relative to VEH-treated hM3DGq mice receiving vehicle (0 mg/kg) only. \**p* < 0.05 and \*\**p* < 0.01, based on Tukey’s post-hoc test following one-way ANOVA (*F*_(3,36)_ = 4.99, *p* < 0.01). All data in (c-e) are means ± SEM with individual values overlaid; *N* = 10 male mice per group and test.

To examine whether elevations in KYNA levels functionally contribute to the disruption of cognitive functions after overactivation of prefrontal astrocytes, we pretreated hM3DGq mice with PF-04859989 2.5 hrs before CNO-mediated activation and 3 hrs before cognitive testing (**Fig. 4c**). PF-04859989 is a brain-penetrable KAT II inhibitor, which efficiently reduces brain KYNA levels after systemic administration^58^. Unlike treatment with a sub-threshold dose (1 mg/kg), administration of PF-04859989 at 10 mg/kg fully restored the deficits in temporal order memory, working memory, and PPI (**Fig. 4d – 4f**) in CNO-treated hM3DGq mice. Importantly, these cognition-enhancing effects were not the result of a general improvement in cognitive performance at baseline conditions. Indeed, when treating non-activated control mice with the same doses of PF-04859989 3 hrs prior to cognitive testing, the drug did not significantly influence temporal order memory, working memory, and PPI (**Supplementary Fig. S5**). Hence, inhibition of KAT II shows a selective efficiency in restoring cognitive impairments in mice with overactivation of prefrontal astrocytes without inducing general pro-cognitive effects under non-activated baseline conditions.

Consistent with these findings, we identified that inhibition of KAT II restores the deficit in neuronal activation of prefrontal PV interneurons after overactivation of prefrontal astrocytes. We pretreated hM3DGq-expressing mice with 0 or 10 mg/kg PF-04859989 2.5 hrs prior to CNO or VEH treatment and 3 hrs prior to the temporal order memory test (**Fig. 5a**). As before, animals were sacrificed 2 hrs after completion of the test and brains were collected for subsequent immunohistochemical analyses of c-Fos and PV (**Fig. 5a**). We found that inhibition of KATII led to a concomitant restoration of the deficits in temporal order memory (**Fig. 5b**) and c-Fos expression in prefrontal PV interneurons **(Fig. 5c**) while sparing the total numbers of PV cells *per se* (**Fig. 5d**).

**Figure 5.**
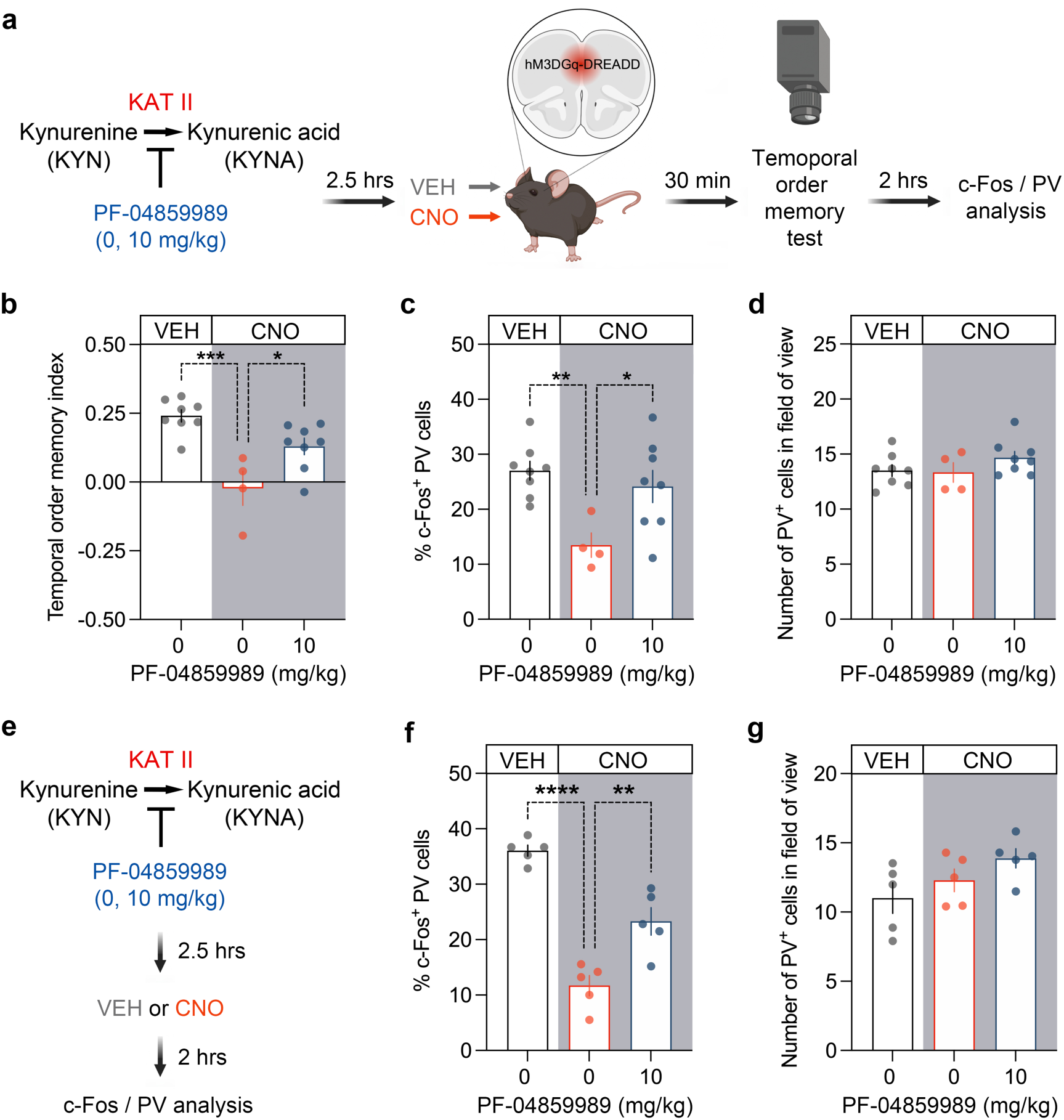
Inhibition of central KYNA production normalizes deficits in PV activation after overactivation of prefrontal astrocytes. **(a)** Schematic representation of the pharmacological rescue study in behaviorally tested mice. Male mice expressing hM3DGq in prefrontal astrocytes were pretreated with the KAT II inhibitor PF-04859989 (0 or 10 mg/kg, i.p.) 2.5 hours before receiving CNO or VEH. The animals were subjected to the temporal order memory test 30 min after VEH or CNO treatment. 2 hours after completion of the test animals were sacrificed and brains were collected for subsequent c-Fox expression analysis. **(b)** Temporal order memory index in the temporal order memory for objects test. \**p* < 0.05 and \*\*\**p* < 0.001, based on Tukey’s post-hoc test following one-way ANOVA (*F*_(2,17)_ = 13.1, *p* < 0.001). **(c)** Percentage of c-Fos positive PV interneurons in behaviorally tested mice. \**p* < 0.05 and \*\**p* < 0.01, based on Tukey’s post-hoc test following one-way ANOVA (*F*_(2,17)_ = 6.075, *p* < 0.01). **(d)** Total number of PV interneurons in behaviorally tested mice. **(e)** Schematic representation of the pharmacological rescue study in behaviorally naïve mice. Male mice expressing hM3DGq in prefrontal astrocytes were pretreated with the KAT II inhibitor, PF-04859989 (0 or 10 mg/kg, i.p.) 2.5 hours before receiving CNO or VEH. 2 hours after treatment the animals were sacrificed, and brains were collected for subsequent c-Fos expression analysis. **(f)** Percentage of c-Fos positive PV interneurons in behaviorally naïve mice. \*\**p* < 0.01 and **** *p* < 0.0001, based on Tukey’s post-hoc test following one-way ANOVA (*F*_(2,12)_ = 42.82, *p* < 0.0001). **(g)** Total number of PV interneurons in behaviorally naïve mice. For (b-d), *N* = 8 (for VEH-0mg/kg PF-04859989 and CNO-10mg/kg PF-04859989) and *N* = 4 (for CNO-0mg/kg PF-04859989) male mice. For (f-g), *N* = 5 male mice per group. All data are means ± SEM with individual values overlaid.

To ensure that these cellular effects are primarily driven by the overactivation of astrocytes and increased KYNA levels rather than by behavioral testing *per se*, we conducted the same immunohistochemical analyses in behaviorally naïve animals (**Fig. 5e**). Consistent with the behaviorally tested animals, overactivation of prefrontal astrocytes significantly decreased c-Fos positive PV interneurons in the PFC. This deficit was mitigated by inhibition of KATII (**Fig. 5f**), in the absence of concomitant effects on the number of PV-positive interneurons *per se* (**Fig. 5g**). Taken together, under conditions of overactivation of prefrontal astrocytes, our findings identify a causal link between increased KYNA levels, impairments in short-term memory and pre-attentive processing, and deficits in the activation of PV interneurons in the PFC.

## DISCUSSION

The present study shows that overactivation of prefrontal astrocytes impairs working memory, temporal order memory, and pre-attentive filtering in both male and female mice and establishes the link to the metabolic pathway of central kynurenines. These findings extend and corroborate previous studies highlighting the importance of astrocytes in numerous behavioral and cognitive domains, including goal-directed behavior^59^, drug seeking and addiction^43, 60^, short-term^61^ and long-term^62^ spatial memory, and motor learning^63^. Our data are in line with the recent study by Delepine et al.^63^, who used the same hM3DGq-DREADDs system in mice to show that chemogenetic activation of cortical astrocytes impairs performance in motor learning and execution tasks. Contrary to these cognition-disrupting effects, however, chemogenetic activation of astrocytes has also been found to induce pro-cognitive effects. For example, using the same hM3DGq-DREADDs system in mice, Adamsky et al.^61^ previously found an enhancement of spatial short-term memory and contextual fear memory recall when astrocytes were chemogenetically activated in the hippocampus. Hence, increasing the functional activity of astrocytes may have beneficial or detrimental effects on behavior and cognition depending on brain region and/or the specificity of the behavioral or cognitive task.

The identification of a pathophysiological link between overactivation of prefrontal astrocytes, increased production of prefrontal KYNA and subsequent disruption of cognition is of particular importance. We found that overactivation of prefrontal astrocytes shifts the KYNA/KYN ratio towards higher KYNA levels in the PFC, whereas it spared other metabolites of the KYN pathway. Importantly, we also showed that pharmacological pretreatment with a KATII inhibitor, which is known to reduce central KYNA levels when administered peripherally^58^, fully restored the deficits in temporal order memory, working memory, and PPI. Taken together, our study provides the first line of evidence suggesting that the disruption of short-term memory and pre-attentive filtering after overactivation of prefrontal astrocytes involves the metabolic pathway of central KYN.

Given its inhibitory actions on NMDA receptors^36, 64^, it is thought that excessive central KYNA impairs cognitive functions by altering prefrontal neurotransmission and activity akin to synthetic NMDA receptor antagonists, such as MK-801^65^, phencyclidine^66^ or ketamine^58^. In support of this hypothesis, studies in rodents show that elevating brain levels of KYNA is sufficient to disrupt various cognitive functions^36^. Of note, increasing brain KYNA levels through systemic administration of KYN leads to behavioral and cognitive alterations that strongly resemble those observed here, including impairments in PPI^38^ and working memory^36, 37, 67^. These phenotypic similarities reinforce our hypothesis suggesting that the increase in KYNA plays a key role in mediating the disruption of short-term memory and sensorimotor gating through overactivation of prefrontal astrocytes.

Our study also sheds light on possible neural mechanisms linking the overactivation of prefrontal astrocytes, alterations in the KYN pathway, and disruption of cognitive functions. Specifically, the c-Fos expression analyses demonstrated an overall increase in neuronal activation after chemogenetic activation of prefrontal astrocytes, but a significant decrease in the activation of PV interneurons in the PFC. The latter effect is consistent with and extends previous findings suggesting that synthetic NMDAR antagonists, such as ketamine, phencyclidine and MK-801, act on prefrontal excitability via reducing the activity of PV cells.^56,68,69-71^ Furthermore, we found that inhibition of KATII prior to prefrontal astrocyte stimulation normalizes the reduction in PV cell activation concomitantly to the restoration of the cognitive impairments. Based on these data and the findings obtained in previous studies,^56,68,69^ we propose that KYNA may readily act on NMDAR expressed by PV interneurons and thereby reduces their activity and disinhibits overall PFC activity.

The identification of elevated prefrontal KYNA levels in our astrocyte DREADDs model also recapitulates the findings of increased cortical and/or cerebrospinal fluid KYNA levels in patients with schizophrenia or bipolar disorder^16, 34, 35, 40, 41^. Moreover, our results are consistent with and extend previous findings providing an association between increased prefrontal KYNA levels and disruption of attention-related cognitive functions in a subgroup of patients with schizophrenia.^16^ While the precise cellular sources of increased KYNA production in schizophrenia and other psychotic disorders remain elusive, our data indicate that an astrocytic origin is both plausible and pathophysiologically relevant. In line with this notion, astrocytic anomalies have been implicated repeatedly in the pathophysiology of both schizophrenia and bipolar disorder. Such anomalies include increased expression of cellular markers indicative of astrocyte activation, such as GFAP^14, 15, 72, 73^, and transcriptomic alterations indicative of increased expression of astrocytic gene sets in cortical post-mortem samples^18-21^. Furthermore, a recent RNA sequencing-based gene enrichment study identified a set of signature genes that were specific to astrocytes and up-regulated in post-mortem brain samples obtained from patients with schizophrenia relative to healthy controls^22^. Taken together, there is accumulating evidence for a molecular basis of astrocytic anomalies in schizophrenia and other psychosis-related disorders, which tentatively point towards an upregulation of astrocytic gene expression in cortical areas of affected individuals. We deem our study a valuable preclinical addition to these findings, as it identifies a causal link between overactivation of prefrontal astrocytes, increased production of KYNA, changes in prefrontal neuronal activity patterns, and disruption of cognitive functions relevant to psychotic disorders and beyond.

The precise molecular mechanisms whereby activation of astrocytes stimulates KYNA production remain to be explored further. Previous attempts to identify these mechanisms have largely focused on the possible involvement of inflammatory stimuli, such as the pro-inflammatory cytokines interleukin (IL)-1β and IL-6, which were both found to stimulate the production of KYNA in human astrocyte cultures^74, 75^. The modulation of enzymes of the KYN pathway is one plausible mechanism by which these cytokines increase KYNA production in astrocytes^74, 75^. Intriguingly, pro-inflammatory cytokines such as IL-1β elevate intracellular Ca^2+^ levels in astrocytes similarly to astrocytic Gq pathway activation^76, 77^. Moreover, a previous study found that the enzymatic activity of KAT enzymes is enhanced by Ca^2+^,^57^ providing a possible mechanism by which elevations in Ca^2+^ can stimulate the conversion of KYN into KYNA. If confirmed by future studies, our findings reported here and by others before^74, 75^ indicate that increasing astrocytic Ca^2+^ signaling in response to activation of the Gq pathway could represent a common mechanism whereby both inflammatory^74, 75^ and non-inflammatory (e.g. activation of hM3DGq) signals stimulate the production and/or release of astrocytic KYNA.

While our data emphasize a critical role of the metabolic pathway of central KYN in mediating the disruption of cognitive functions after overactivation of prefrontal astrocytes, there are alternative (but not mutually exclusive) mechanisms whereby activation of the Gq pathway in cortical astrocytes could affect behavior and cognition. For example, using the same hM3DGq-DREADD system in mice, Delepine et al.^63^ recently found that overactivation of astrocytes upregulates the expression of the glycine transporter 1 (GlyT1). GlyT1 is expressed on astrocytes and mediates clearance of extra-cellular glycine in the proximity of NMDAR-containing synapses^78^. Because glycine is an obligatory, allosteric co-agonist for NMDA receptor activation upon stimulation by glutamate^79^, increased or reduced expression of GlyT1 is expected to blunt or boost NMDA receptor functions, respectively^80^. Interestingly, KYNA inhibits NMDA receptor functions by binding to the receptor’s glycine co-agonist site and at higher levels to the glutamate binding site^64^. Thus, the functional interference with the glycine co-agonist activity may reflect a common pathway by which functional overactivation of prefrontal astrocytes modulates cognitive functions, be it mediated by reducing the availability of extracellular glycine through GlyT1 upregulation or by blocking the glycine co-agonist site through KYNA. In line with this notion, pharmacological inhibition of GlyT1 and enzymes stimulating the production of KYNA (e.g. KAT II) have been shown to produce similar pro-cognitive effects^58, 80, 81^. Whether or not the cortical KYN pathway and/or cortical GlyT1 offer targets for cognitive enhancement in psychiatric disorders and beyond remains a matter of ongoing discussion and research^67, 82-84^.

We acknowledge a number of limitations in our study. First, some of the experiments presented here, including the postmortem evaluation of c-Fos expression, measurements of the metabolites of the KYN pathway, and pharmacological interventions with the KATII inhibitor were conducted in male animals expressing hM3DGq only. Therefore, we cannot exclude possible sex-dependent outcomes in these tests. However, the initial characterization of the behavioral and cognitive effects of prefrontal astrocyte activation revealed highly comparable outcomes in male and female mice. In keeping with these sex-independent effects, we deemed it appropriate to conduct all subsequent experiments using animals of one sex only in order to minimize the number of animals. Second, KYNA levels were measured in PFC bulk tissue, which did not enable us to differentiate between intracellular and extracellular KYNA levels. Third, because the target area of primary interest was the PFC in our study, our findings do not allow us to generalize our findings to other brain regions. Thus, the extent to which KYNA could modulate behavior and cognition after astrocytic activation in other brain regions remains to be investigated.

Notwithstanding these limitations, our study provides the first causal link between overactivation of prefrontal astrocytes, a shift in the KYNA/KYN ratio towards higher KYNA levels, disruption of cognitive functions, and alterations in neuronal activation patterns. Our findings are especially relevant for psychosis-related diseases such as schizophrenia and bipolar disorder, which involve PFC-associated cognitive impairments, increased cortical expression of astrocytic markers^19, 85^, alterations in prefrontal PV interneuron signaling^19, 31, 32^, and increased central production of KYNA^36, 40^.

## MATERIAL AND METHODS

### Animals

All experiments were performed using male or female C57BL6/N mice (Charles Rivers, Sulzfeld, Germany). They were group-housed (4-5 animals per cage) in individually ventilated cages (Allentown Inc., Bussy-Saint-Georges, France). The cages were kept in a specific-pathogen-free (SPF) holding room, which was temperature- and humidity-controlled (21 ± 3 °C, 50 ± 10%) under a reversed light–dark cycle (lights off: 09:00 AM–09.00 PM). All animals had *ad libitum* access to standard rodent chow (Kliba 3336, Kaiseraugst, Switzerland) and water throughout the entire study. All procedures were conducted during the dark cycle and had been previously approved by the Cantonal Veterinarian’s Office of Zurich. All efforts were made to minimize the number of animals used and their suffering. An overview of the different cohorts of animals and the respective numbers used are provided in the **Supplementary Table 1**. In addition, the number of animals used in each experiment is specified in the legends of the main and extended figures.

### DREADD system

The DREADD system was based on a recombinant adeno-associated virus serotype 9 (AAV9) that expresses hM3DGq under an astrocyte-specific promoter (hgfaABC1D), encompassing a fluorescent mCherry tag (AAV9-hgfaABC1D-hM3DGq-mCherry; **Fig. 1a**). Some experiments involved an EGFP-tagged control virus (ConV) with the same promoter (AAV9-hgfaABC1D-EGFP) and the GCaMP6s Ca^2+^ sensor (AAV9-hgfaABC1D-GCaMP6s). All AAVs were produced and purchased from the Viral Vector Facility of the University of Zurich, Switzerland (www.vvf.uzh.ch) and were injected into the PFC using bilateral stereotaxic injections, as described in the **Supplementary Information**. hM3DGq was activated with 1 mg/kg clozapine-N-oxide (CNO, BML-NS105-0025, Enzo Life Sciences, Switzerland) dissolved in 0.9% NaCl (B. Braun, Switzerland). The dose of 1 mg/kg was chosen based on previous chemogenetic studies in rodents^46, 53, 86, 87^. Treatment with the vehicle (VEH; 0.9% NaCl) served as control treatment in hM3DGq-expressing mice. ConV-expressing mice receiving CNO or VEH were used to exclude non-selective effects of CNO^52^.

For all experiments, CNO (1 mg/kg) or VEH were given via the micropipette-guided drug administration (MDA) method, a non-invasive oral administration technique described in detail elsewhere^46, 47^.

### In vivo two-photon Ca^2+^ imaging of prefrontal astrocytes in awake mice using microprisms

In vivo Ca^2+^ imaging using two-photon microscopy in awake mice was applied to ascertain the effectiveness and temporal dynamics of the hM3DGq-DREADD system in prefrontal astrocytes. To this end, we unilaterally co-injected the hM3DGq construct with a viral construct expressing the Ca^2+^ sensor, GCaMP6s, into the PFC of adult mice. To access the PFC, a right-angled microprism attached to a cranial window was implanted into the subdural space within the fissure^45^ (**Fig. 1e** and **Supplementary Information**). One week after surgery the animals were trained for the awake in vivo two-photon Ca^2+^ imaging for a total period of 2 weeks. Imaging was then performed at baseline and after administration of either VEH or CNO via MDA. Methodological details regarding the surgical procedure and training of mice, as well as the in vivo two-photon imaging experiments are available in the **Supplementary Information**.

### Behavioral and cognitive testing

Behavioral and cognitive testing included tests for innate anxiety-like behavior and exploratory activity (light-dark box and open field tests), a temporal order memory test for objects, a spontaneous alternation task for working memory in the Y-maze, and a prepulse inhibition (PPI) test of the acoustic startle reflex for pre-attentive filtering. A detailed description of the methodological procedures and rationale of inclusion for each behavioral and cognitive test are provided in the **Supplementary Information**.

### Immunohistochemistry and laser-scanning confocal microscopy

Immunofluorescent staining and laser-scanning confocal microscopy were used to ascertain the cellular expression pattern of the hM3DGq construct and to analyze neuronal activation in response to overactivation of prefrontal astrocytes. The latter was conducted using quantification of c-Fos expression, which was assessed 2 hrs after CNO or VEH administration in behaviorally naïve animals, or 2 hrs after completion of the temporal order memory test (behaviorally exposed animals). Methodological details regarding immunohistochemistry, image acquisition, and analyses are available in the **Supplementary Information**.

### Quantification of brain metabolites of the kynurenine pathway

Samples of PFC bulk tissue (including anterior cingulate, prelimbic, and infralimbic cortices) were collected and processed as described in the **Supplementary Information**. Brain metabolites of the KYN pathway were quantified using liquid chromatography-nanoelectrospray ionization tandem mass spectrometry as described in the **Supplementary Information**. Besides KYN and KYNA, the following metabolites were quantified as well: tryptophan (TRP), quinolinic acid (QUIN), and 3 hydroxykynurenine (3-HK).

### Kynurenine aminotransferase II inhibition

Kynurenine aminotransferase II (KATII) was inhibited with 1 mg/kg or 10 mg/kg PF-04859989 hydrochloride (PZ0250, Sigma-Aldrich), which was dissolved in sterile water and freshly prepared prior to each experiment. For animals subjected to behavioral testing, 1 or 10 mg/kg PF-04859989, or sterile water (vehicle, 0 mg/kg) only, were injected i.p. using an injection volume of 5 mL/kg 3 hrs prior to each behavioral test. For postmortem analyses of c-Fos expression, sterile water (vehicle) or 10 mg/kg PF-04859989 was administered either 5 hours (behaviorally naïve animals) or 8 hrs (behaviorally exposed animals) prior to tissue collection. The doses and post-injection interval were chosen based on previous dose-response studies in rodents^58, 88^.

### Statistical Analyses

All statistical analyses were performed using SPSS Statistics (version 29.0, IBM, Armonk, NY, USA) and Prism (version 9.0; GraphPad Software, La Jolla, CA, USA). Statistical significance was set at *P* < 0.05. Detailed information regarding the statistical analyses used for each experiment is available in the **Supplementary Information**.

## Acknowledgments

We thank Endre Laczko and Kurt Stefan Schauer from the Functional Genomics Center Zurich (FGCZ), Switzerland, for their technical assistance in liquid chromatography-nanoelectrospray ionization tandem mass spectrometry. This work was financially supported by the Swiss National Science Foundation (grant No. PZ00P3_202149 and grant No. P2ZHP3_174868 awarded to T.N.; grant No. 310030_188524, awarded to U.M.). Additional financial support was provided by the Brain & Behavior Research Foundation (grant No. 30963 awarded to T.N.).

## Contributions

V.B., J.F., S.M.S., U.W., and M.W. were involved in the acquisition, analysis, and interpretation of the study data; T.N. and U.M. were involved in the conception and design of the study and analysis and interpretation of the study data; T.N., A.S., B.W., and U.M. supervised research; T.N. and U.M. wrote the initial manuscript draft; all authors contributed to the reviewing and editing of the manuscript, and have given final approval for the version to be published.

## Ethics declaration

All authors declare no competing interests. The funders of the study had no role in the study design, data collection and analysis, decision to publish, or preparation of the manuscript.

## Data availability

Source data are provided with this paper. Any additional data that we inadvertently missed will be shared upon reasonable request.

