## Supplementary information for "Overactivation of prefrontal astrocytes impairs cognition through the metabolic pathway of central kynurenines"

### SUPPLEMENTARY METHODS

#### Animals

| Cohort | Experimental procedure | Experimental groups | Sample size |
|---|---|---|---|
| 1 | Awake in vivo two-photon imaging | | 4 |
| 2 | Expression pattern of hM3DGq | | 10 |
| 3 | Behavioral testing of hM3DGq-expressing male mice | VEH<br>CNO | 10<br>10 |
| 4 | Behavioral testing of ConV-expressing male mice | VEH<br>CNO | 10<br>10 |
| 5 | Behavioral testing of hM3DGq-expressing female mice | VEH<br>CNO | 10<br>10 |
| 6 | c-Fos expression analyses of behaviorally exposed hM3DGq-expressing male mice | VEH<br>CNO | 8<br>8 |
| 7 | Kynurenine metabolite measurement in hM3DGq-expressing male mice | VEH<br>CNO | 10<br>10 |
| 8 | Behavioral testing of hM3DGq-expressing mice after pharmacological inhibition of KATII | vehicle/VEH<br>vehicle/CNO<br>1mg/kg PF-04859989/CNO<br>10mg/kg PF-04859989/CNO | 10<br>10<br>10<br>10 |
| 9 | Behavioral testing of C56BL6/N male mice after pharmacological inhibition of KATII | vehicle<br>1mg/kg PF-04859989<br>10mg/kg PF-04859989 | 9<br>10<br>10 |
| 10 | c-Fos expression analysis of behaviorally tested hM3DGq-expressing male mice after pharmacological inhibition of KATII | VEH<br>CNO<br>10mg/kg PF-04859989/CNO | 8<br>4<br>8 |
| 11 | c-Fos expression analysis of behaviorally naïve hM3DGq-expressing male mice with pharmacological inhibition of KATII | VEH<br>CNO<br>10mg/kg PF-04859989/CNO | 5<br>5<br>5 |

**Supplementary Table 1:** Overview of the different cohorts of animals and sample sizes used in this study.

#### AAV Constructs

The following recombinant adeno-associated viruses (AAVs) were used: AAV9-GFAP-hM3DGq-mCherry (hM3DGq; physical titre: 9.2 × 10E11 vg/mL), AAV9-GFAP-EGFP (ConV; physical titre: 7.6 × 10E11 vg/mL), AAV9-GFAP-GCaMP6s (GCaMP6s; physical titre: 3.2 × 10E12 vg/mL). The hgfaABC1D GFAP promoter was used for all constructs to ensure astrocyte-specific expression. All

AAVs were produced and purchased from the Viral Vector Facility of the University of Zurich, Switzerland (www.vvf.uzh.ch).

**Stereotaxic Surgery**

The stereotaxic surgery was performed when the animals reached 10 weeks of age, using methods established and validated before[1, 2]. Anesthesia of the animals was induced by inhalation of 4% isoflurane (ZDG9623V, Baxter, Switzerland) in oxygen. After anesthesia induction, the heads of the animals were shaved, and vitamin A cream (Bausch & Lomb Swiss AG) was applied to the eyes to avoid dehydration. The animals were injected with the analgesic Temgesic [buprenorphine (0.1 mg/kg, s.c.), Reckitt Benckiser, Switzerland] and fixed into the stereotaxic frame (MTM-3, World Precision Instruments, USA) while kept under constant isoflurane/oxygen flow [1 to 3% isoflurane in oxygen (600 ml/min)]. All animals were kept on a temperature-controlled heating plate (ATC1000, World Precision Instruments, USA) during the entire surgical procedure to avoid anesthesia-induced hypothermia. Prior to incision, local anesthetics was administered subcutaneously at the incision site (50 µl of a mixture of 1 mL lidocaine (10mg/mL) and 1 mL bupivacaine (5 mg/mL) in 2 mL saline). A longitudinal incision of the skin was made to expose the skull. The skull was cleaned from connective tissue, and the bone, above the target area, was removed using a micro drill (OmniDrill35, World Precision Instruments, USA) with a rose burr (ø 0.3 mm). Intracerebral injections were performed using a NanoFil needle and syringe (NANOFIL, NF35BV, World Precision Instruments, USA) connected to an automated pump (UMP3T-1, World Precision Instruments, USA).

Mice received a single, bilateral injection of either the AAV9-hGFAP-hM3DGq-mCherry or AAV9-hGFAP-EGFP into the medial portion of the prefrontal cortex (anteroposterior [AP] = +1.8 mm, mediolateral [ML] = +0.3 mm, dorsoventral [DV]= −1.9 mm). Solutions were injected at an infusion rate of 10 nL/s and a total volume of 450 nL. After insertion of the needle, a small drop of Histoacryl (B. Braun, Switzerland) was applied at the site of injection to avoid reflux of the injected substances. After injection, the needle was kept in place for 5 min to avoid reflux of the substances before retracting it. Incisions were sutured with a surgical thread (G0932078, B. Braun, Switzerland), and the animals were placed in a temperature-controlled chamber (Harvard Apparatus, USA) until full recovery from anesthesia. After the surgery, the animals were placed back in their home cage and closely monitored for three consecutive days after surgery.

**DREADD Activation**

hM3DGq was activated with 1 mg/kg clozapine-N-oxide (CNO, BML-NS105-0025, Enzo Life Sciences, Switzerland) dissolved in 0.9% NaCl (B. Braun, Switzerland). The dose of 1 mg/kg was chosen based on previous chemogenetic studies in rodents[1, 3-5]. CNO (1 mg/kg) or vehicle (0.9% sterile saline, VEH) were given via the micropipette-guided drug administration (MDA) method as described in detail elsewhere[5, 6]. For behavioral testing and tissue collection for brain metabolite measurements, CNO or VEH were given 30 min prior to testing or tissue collection. For c-Fos analyses, CNO or VEH were given either 2 hrs (behaviorally naïve cohort) or 5 hrs (behaviorally tested cohorts) prior to tissue collection. In the awake two photon imaging experiments, VEH or CNO were given during imaging sessions.

**Immunohistochemistry**

The animals were deeply anesthetized with an overdose of pentobarbital (Esconarkon ad us. vet., Streuli Pharma AG, Switzerland) and transcardially perfused with ice-cold artificial cerebrospinal fluid (pH 7.4)[1, 7, 8]. The brains were immediately removed from the skull and postfixed in 4% PFA for 6 hours before cryoprotection in 30% sucrose in PBS for 24-48 hours. The brains were cut coronally with a sliding microtome at 30 μm (eight serial sections) and stored at −20°C in cryoprotectant solution [50 mM sodium phosphate buffer (pH 7.4) containing 15% glucose and 30% ethylene glycol; Sigma-Aldrich, Switzerland] until further processing.

Immunofluorescent stainings were performed according to previously established protocols[1, 5, 7, 8]. Briefly, the brain sections were rinsed in tris buffer (pH 7.4) before incubating with primary antibodies (mCherry, rat monoclonal (16D7), M11217, Invitrogen, Switzerland, 1:1000; s100β, rabbit monoclonal (EP1576Y), ab52642, Abcam, Switzerland, 1:1000; Iba1, rabbit polyclonal, 019-19741, Wako Chemicals, USA, 1:2000; NeuN, rabbit monoclonal (EPR12763), ab177487, Abcam, Switzerland, 1:500; c-Fos, rabbit, monoclonal (9F6), 2250, Cell Signaling, USA, 1:1000; PV, guinea pig, polyclonal, 195 004, Synaptic Systems, Germany, 1:1000). The primary antibodies were diluted in tris buffer containing 0.2% Triton X-100 and 2% normal serum. The sections were incubated free-floating under constant agitation (100 rpm) overnight at 4°C. The following day, sections were washed three times for 10 min in tris buffer before a 30-min incubation period with secondary antibodies (Alexa488 (Molecular Probes, Eugene, USA, 1:1000), Cy3 (Jackson ImmunoResearch, UK, 1:500), or Cy5 (Jackson ImmunoResearch, UK, 1:500), and DAPI (1 mg/mL $H_2O$; Thermo Fisher Scientific, Switzerland, 1:3000)) diluted in tris buffer containing 2% normal serum at room temperature. After incubation, which was shielded from light, the sections were washed 3 × 10 min in tris buffer, mounted onto gelatinized glass slides, coverslipped with Dako fluorescence mounting medium (S3023, Agilent, Switzerland), and stored in the dark at 4°C until image acquisition.

Immunofluorescence images were captured by laser scanning confocal microscopy or with Airyscan confocal microscopy (Zeiss LSM800 with Airyscan). For assessing the selectivity of DREADD construct expression, 6 randomly selected z-stacks within the area of construct expression were acquired per animal using a 40× (oil, NA 1.4) objective with a zoom of 0.45. Optical sections were acquired with a resolution of 1024 x 1024 pixels per section. Higher resolution image stacks for representative images of cell type specificity were acquired in Airyscan mode using a 40× lens, NA 1.4, oil and processed using the default settings provided by ZEN 2.6 blue edition software (Zeiss, Switzerland). For c-Fos expression analysis, tile scans were acquired using 10× lens, NA 0.45, air and processed using the stitching tool provided by Zen 2.6 blue edition software (Zeiss, Switzerland). For c-Fos/PV co-localization analysis, 6 randomly selected images within the area of construct expression were acquired per animal using a 25× (oil, NA 0.8) objective with a zoom of 1. Images were acquired with a resolution of 1024 x 1024 pixels per section. Imaging of the individual cohorts was conducted on the same experimental day and imaging settings were kept constant throughout an entire imaging day. Final illustrations were rendered in ImageJ or Imaris, where contrast and brightness were adjusted across an entire image.

Image analysis was performed using the ImageJ software by an experimenter blinded to the experimental conditions. For assessing the selectivity of DREADD construct expression, the total number of mCherry$^+$, s100β$^+$, NeuN$^+$, mCherry$^+$/s100β$^+$, and mCherry$^+$/NeuN$^+$ cells were counted within each image. Expression selectivity was then calculated by dividing the number of colocalized cells with the number of total mCherry$^+$ cells and multiplied by 100. For cFos expression analysis, 3 tile scans containing the PFC were acquired per animal. The region of interests (ROI) was drawn around the PFC using the polygon selection tool as outlined in main Fig 1b. Gaussian filter, background subtraction and a threshold were applied to the stitched images. The settings were kept constant throughout the entire analysis. The number of cells within the area of the ROI was calculated using the analyze particle plugin. For c-Fos/PV colocalization analysis, the total number of c-Fos$^+$, PV$^+$, and c-Fos$^+$ / PV$^+$ cells were counted manually within each image. The percentage of c-Fos$^+$ / PV$^+$ cells were calculated by dividing the number of colocalized cells with the number of total PV+ cells and multiplied by 100.

**In Vivo Two-Photon Imaging**

*Microprism cranial window assembly:* On the day of the surgery, a custom-made right angle microprism (1.5-mm side length and 1-mm width, S-BSL7, protected aluminum coating on hypotenuse surface to enable internal reflection; Optosigma) was bonded to a circular glass window (3 mm diameter coverslip) using UV curing optical adhesive (Norland #81).

*Surgery and virus injection:* Surgery and AAV injections were performed using methods established and validated before[9, 10]. Headplate implantation, craniotomy, AAV injection and microprism implantation were performed under midazolam (5 mg/kg), fentanyl (0.05 mg/kg), and medetomidine (0.5 mg/kg) anesthesia. First, a chrome steel head plate was implanted. In brief, anesthetized animals were fixed in a stereotaxic frame and an incision was made along the midline to expose the skull. The bone was cleaned, and a bonding agent (Prime & Bond) applied to the skull and polymerized with blue light. A round head plate was attached to the exposed bone using light-cured dental cement (Tetric EvoFlow). Next, a craniotomy was performed over the PFC using a dental drill (rotate, H-4-002). The skull was first thinned and carefully removed in small bone fragments leaving the dura intact. Next, a pipette and hydraulic pump were used to unilaterally inject a 1:1 mixture of AAV9-GFAP-hM3DGq-mCherry and AAV9-GFAP-GCaMP6s virus (injection volume of 600 nL and injection speed of 10 nL/s) into the PFC (1.2 mm below cortical surface) opposing the site of microprism implantation. Next, the microprism was implanted, whereby an incision in the dura along the side of the sinus was created where the microprism was inserted. The microprism was then gently inserted into the subdural space within the fissure so that the prism surface sat flush opposite the hemisphere to be imaged (with the cerebral falx between the microprism surface and PFC cortex to be imaged). The area beneath the microprism (i.e., the medial portion of the PFC of the contralateral site, which was not imaged) was compressed but remained intact. A rim of dental cement was then used to secure the glass cover slip in place.

*Training for awake in vivo two-photon imaging:* One week after surgery, the training of animals for awake imaging commenced. During the first five days, mice were handled during which their head was manually fixed by holding the head plate by the experimenter. During these five days, the handling and manual fixation times were gradually increased from 5 min handling and no fixation (first day) to 20 min handling and 5 min fixation (fifth day). At the end of each session, a reward in the form of 40% condensed milk was given via MDA. On day 6, the animals were introduced to being head fixed in the awake imaging setup, whereby the animals were allowed to explore the imaging setup with manual fixation by the experimenter. As of day 7, the animals were being gradually head-fixed in the imaging setup. As for the handling and manual fixation, the times during which animals were head fixed to the imaging setup was gradually increased from 5 to 45 min. The animals were being rewarded with 40% condensed milk via MDA during head fixation and after each session.

*In Vivo Two-Photon Imaging:* Imaging commenced 3 weeks after virus injection and microprism implantation using a custom-built two-photon laser-scanning microscope[11]. A 16× water immersion microscope objective was used (W Plan-Apochromat 16×/1.0 DIC VIS-IR, Zeiss). GCaMP6s and mCherry were excited at 940 nm and 1100 nm with a Ti:sapphire laser (Mai Tai; Spectra-Physics) with power between 10 and 30 mW. Fluorescence emission was detected with a GaAsP photomultiplier module (Hamamatsu Photonics) fitted with 520/550 nm band pass filter or a 607/670 band pass filter and separated by a 560 nm dichroic mirror (BrightLine; Semrock). The two-photon laser-scanning microscope was controlled by a customized version of “ScanImage” (r3.8.1; Janelia Research Campus). All imaging was performed in head-fixed awake mice on an air-lifted platform.

High resolution images (512 x 512 pixels) at a frequency of 0.74 frames per second and averages of 20 frames of every spot were acquired at the start of every imaging session to ensure localization of the imaging spot over multiple sessions. For $Ca^{2+}$ response measurements, images (128 x 128 pixels) were acquired at a frequency of 5.92 frames per second without averaging. For the first imaging session, 10 min baseline activity was recorded. After 10 min, the imaging session was briefly stopped to administer VEH via MDA. Imaging was continued immediately after administration (~10 to 30 s) and activity after VEH was recorded for 15 min. After this, the session was briefly stopped again and CNO (1mg/kg) was administered via MDA and imaging was continued for 30 min thereafter. One hr after termination of the CNO imaging session, the animals were reimaged for 5 min to assess the duration of CNO-induced overactivation of prefrontal astrocytes. For the second imaging session (at least 48 hrs after the first session), 10 min baseline activity was recorded before administration of CNO (1 mg/kg) as described above. Imaging after CNO was continued for 30 min.

*Quantification and analysis:* Image analysis was performed using the ImageJ software. For each field of view, baseline F0 was defined as the average pixel intensity during the baseline activity recordings of the respective session. This baseline was then subtracted from all pixel intensities for the remaining imaging session. The resulting difference was divided by F0 to obtain dF/F0. To correct for intensity changes due to movement and bleaching, dF/F0 of the GCaMP6f signal was normalized to dF/F0 of the mCherry signal for each frame.

**Kynurenine Aminotransferase II Inhibition**

Kynurenine aminotransferase II (KATII) was inhibited with 1 mg/kg or 10 mg/kg PF-04859989 hydrochloride (PZ0250, Sigma-Aldrich), which was dissolved in sterile water and freshly prepared prior to each experiment. For animals subjected to behavioral testing, 1 or 10 mg/kg PF-04859989, or sterile water (vehicle, 0 mg/kg) only, were injected i.p. using an injection volume of 5 mL/kg 3 hrs prior to each behavioral test. For postmortem analyses of cFos expression, sterile water (vehicle, 0 mg/kg) or 10 mg/kg PF-04859989 was administered either 5 hrs (behaviorally naïve animals) or 8 hrs (behaviorally tested animals) prior to tissue collection. The doses and post injection interval were chosen based on previous studies in rodents[12, 13].

**Light-Dark Box Test**

A light-dark box test was used to measure innate anxiety-like behavior[14]. The apparatus consisted of four identical multi-conditioning boxes (Multi Conditioning System, TSE Systems, Germany), each containing a dark (1 lux) and a bright (100 lux) compartment. The two compartments were separated from each other by a dark plexiglass wall with an integrated, electrically controlled door. To start a trial, each mouse was placed in the dark compartment. After 5 s, the door automatically opened, allowing access to both the dark and bright compartments for 10 min. The measurements collected from this test included the distance moved and time spent in the light compartment.

**Open Field Test**

A standard open field exploration task served to assess spontaneous locomotor activity and innate anxiety-like behavior[14]. The apparatus consisted of four identical open-field arenas [40 cm (length) by 40 cm (width) by 35 cm (height)] made of white polyvinyl chloride (OCB Systems Ltd., UK). It was positioned in a testing room with diffused lighting (~30 lux in the center of the arena). A digital camera was mounted above the arena, captured images at a rate of 5 Hz, and transmitted them to a PC running the EthoVision (Noldus Technology, The Netherlands) tracking system. The animals were recorded for 25 min before they were placed back into their home cage. For the purpose of data collection, the arena was conceptually partitioned into two areas: a center zone (measuring 10 cm by 10 cm) in the middle of the area and a peripheral zone occupying the remaining area. The measurements collected from this test included the total distance moved, and the distance moved in the center zone.

**Temporal Order Memory Test**

A temporal order memory test for objects was used to the animals' capacity to discriminate the relative recency of stimuli[15]. The test apparatus consisted of an open field as described above, with minor modifications (see below). The test procedure consisted of three consecutive phases, which were each separated by 60 min.

*Sample phase 1:* For this phase, a first pair of identical objects (blue aluminum hairspray bottles, 250 ml, 20 cm high) were placed in the open-field arena in opposing corners approximately 5 cm from the walls. To start a trial, the animals were gently placed into the center of the open field and were allowed to freely explore the objects for 10 min. They were then removed from the apparatus again and kept in a holding room for 60 min before the start of the next phase.

*Sample phase 2:* For this phase, a novel pair of identical objects (LEGO Duplo brick pile, 15 cm high) were placed in the open-field arena, thereby allocating them in the same position as the first pair of objects (see above). To start a trial, the animals were gently placed into the center of the open field again and were allowed to freely explore the novel pair of objects for 10 min. They were then removed from the apparatus once more and kept in a holding room for 60 min before the start of the actual test phase.

*Test phase:* In the test phase, the open field was equipped with one object used in sample phase 1 (temporally more remote object) and one object in sample phase 2 (temporally more recent object), with the corner allocation of the objects being counterbalanced across groups. To start the test trial, the animals were placed into the center of the open field and were allowed to freely explore either object for 5 min. For each animal, a temporal order memory index was calculated by the following formula: [(time spent with phase 1 object)/(time spent with phase 1 object + time spent with phase 2 object)] − 0.5. The temporal order memory index was used to compare the animals' capacity to discriminate the relative recency of stimuli,[15] with values > 0 signifying a capacity to discriminate between the temporally more remote object presented in sample phase 1 and the temporally more recent object presented in sample phase 2. In addition, the relative amount of time exploring the objects in sample phases 1 and 2 of the test was analyzed to measure object exploration per se and to explore possible side preferences while exploring the objects (data not shown).

**Short-term Memory Test in the Y-Maze**

A spontaneous alternation task in the Y-maze was used to assess working memory[16, 17]. This task is based on the innate tendency of rodents to explore novel environments, that is, their preference to investigate a new arm of the Y-maze rather than returning to one that was previously visited[16, 17]. The apparatus was made of transparent Plexiglas and consisted of three identical arms (50 cm × 9 cm; length × width) surrounded by transparent Plexiglas walls 10 cm in height. The three arms radiated from a central triangle (8 cm on each side) and were spaced 120° from each other. The maze was elevated 90 cm above the floor and was positioned in a dimly lit room. A digital camera was mounted above the Y-maze apparatus. Images were captured at a rate of 5 Hz and transmitted to a PC running the EthoVision tracking system (Noldus Information Technology), which calculated the total distance moved (m) in the Y-maze.

To start the test, the animals were gently placed in the center of the Y maze and allowed to explore freely for 5 min, whereby the number and sequence of arm entries (defined as entry of the whole body into an arm) were observed and recorded by an experimenter who was blinded to the treatment conditions. Alternation was defined as entry into the three arms in any non-repeating order (for example, ABC, BAC, CBA). Working memory was indexed by the percentage alternation score, which was calculated as the total number of alternations divided by the possible alternations given the number of arm entries (total number of arm entries - 2). In addition to the analysis of percentage alternation, the total distance moved was recorded and analyzed to assess general activity during the 5-min testing period.

**Prepulse Inhibition of the Acoustic Startle Reflex**

Pre-attentive filtering was assessed using the paradigm of prepulse inhibition (PPI) of the acoustic startle reflex. PPI of the acoustic startle reflex refers to the reduction in startle reaction in response to a startle-eliciting pulse stimulus when it is shortly preceded by a weak prepulse stimulus. The apparatus consisted of four startle chambers for mice (San Diego Instruments, USA) and has been fully described elsewhere[18]. In the demonstration of PPI, the animals were presented with a series of discrete trials comprising a mixture of 4 trial types. These included pulse-alone trials, prepulse-plus-pulse trials, prepulse-alone trials, and no-stimulus trials in which no discrete stimulus other than the constant background noise was presented. The pulse and prepulse stimuli used were in the form of a sudden elevation in broadband white noise level (sustaining for 40 and 20 ms, respectively) from the background (65 $dB_A$), with a rise time of 0.2–1.0 ms. In all trials, 3 different intensities of pulse

(100, 110, and 120 $dB_A$) and 3 intensities of prepulse (71, 77, and 83 $dB_A$) were used. The stimulus-onset asynchrony of the prepulse and pulse stimuli on all prepulse-plus-pulse trials was 100 ms (onset-to-onset).

The protocol used for the PPI test was extensively validated before[8, 19, 20]. A session began with the animals being placed into the Plexiglas enclosure. They were acclimatized to the apparatus for 2 min before the first trial began. The first 6 trials consisted of 6 startle-alone trials; such trials served to habituate and stabilize the animals' startle response and were not included in the analysis. Subsequently, the animals were presented with 10 blocks of discrete test trials. Each block consisted of the following: three pulse-alone trials (100, 110, or 120 $dB_A$), 3 prepulse-alone trials 71, 77, and 83 $dB_A$), 9 possible combinations of prepulse-plus-pulse trials (3 levels of pulse × 3 levels of prepulse), and one no stimulus trial. The 16 discrete trials within each block were presented in a pseudorandom order, with a variable interval of 15 s on average (ranging from 10 to 20 s). For each of the 3 pulse intensities (100, 110, or 120 $dB_A$), PPI was indexed by percent inhibition of the startle response obtained in the pulse-alone trials by the following expression: 100% × [1 − (mean reactivity on prepulse-plus-pulse trials/mean reactivity on pulse-alone trials)], for each animal, and at each of the three possible prepulse intensities. In addition to PPI, reactivity to pulse-alone trials and prepulse-alone trials were also analyzed.

**Liquid Chromatography-Nanoelectrospray Ionization Tandem Mass Spectrometry**

Brain metabolites of the kynurenine pathway were measured using liquid chromatography-nanoelectrospray ionization tandem mass spectrometry (LC-NSI-MS/MS) performed at the Functional Genomic Center Zurich (Metabolomics at FGCZ, University of Zurich).

*Sample collection:* Male mice expressing hM3DGq in the mPFC were treated orally with 1mg/kg CNO or 0.9% saline. 30 min after treatment they were deeply anesthetized with an overdose of pentobarbital (Esconarkon ad us. vet., Streuli Pharma AG, Switzerland) and transcardially perfused with ice-cold artificial cerebrospinal fluid (pH 7.4)[1, 7, 8] to flush out blood derived metabolites of the kynurenine pathway. The brains were immediately removed from the skull and the mPFC was dissected on ice as described before[1, 2, 16]. The mPFC samples from two mice were pooled, weighed and homogenized in 200 µL 80% ice cold methanol using a sterile BioMasher (9791A, Takara, France). The samples were frozen with dry ice and transported to the Functional Genomics Center Zurich, University of Zurich for targeted metabolomic analysis.

*Sample processing:* The thawed samples were centrifuged (15 min / 14 krpm / +4 °C). From each clear, colorless supernatant 180 µL were transferred to 1.5 mL eppendorf tubes, evaporated to dryness under a gentle stream of nitrogen and re-dissolved in 200 µL water containing 10 mM ammonium formate (pH 4.2) and 0.1 % (v/v) formic acid prior to mass spectrometry analysis.

*LC-NSI-MS/MS analysis*: LC-NSI-MS/MS was performed on a TSQ Quantiva triple quadrupole mass spectrometer (ThermoFisher Scientific, United States) coupled to an ACQUITY UPLC M-Class (Waters, United States) system using nanoelectrospray ionization. The analysis was conducted using a capillary column (150 µm ID, 5.5 cm length, 15 µm orifice) created by hand packing a commercially available fused-silica emitter (MSWIL, Netherlands) with HSS T3 separation media (Waters, United States). The mobile phase consisted of 10 mM ammonium formate (pH 4.2) and 0.1 % (v/v) formic acid in water (A1) and 0.1 % (v/v) formic acid in acetonitrile (B1). A 1 µL injection loop was used and the sample (1 µL) was loaded onto the capillary column with 2 µL/min flow at the initial conditions (92.5 % A1, 7.5 % B1) and eluted under isocratic conditions at a flow rate of 2 µL/min over 10 min, following by ramping to 98 % B1 within 1 min and holding at this composition for an additional 4 min. The column was then re-equilibrated at the initial conditions for 5 min before the next injection. The nanoelectrospray source was operated in positive ion mode and the voltage set at 2.7 kV. The ion transfer tube temperature was 250 °C and the radio frequency (RF) lens set at 90 V. The collision gas was Ar at 1.5 mTorr with collision energy of 14 eV and the quadrupoles were operated at a resolution of 0.4 Da for Q1 and of 0.7 Da for Q3. The mass transitions for monitoring the analytes were m/z 190 → 144.1 for KYNA, 209.1 → 192.2 for KYN, m/z 205.1 → 146.1 for TRP, m/z 168 → 124 for QUIN, and m/z 225 → 162.1 for 3-HK, respectively.

The quantitation of the analytes was done using the mass spectrometers vendor software package Quan Browser in the software suit Xcalibur based on the peak areas and the constructed calibration curves. Calibration curves were constructed for each analyte during each analysis using a series of standard solutions of analytes. Pmole/mg tissue was calculated using the following calculation: ((nM of metabolite measured in injected volume) * 0.001 * 200)/(tissue weight).

**Statistical Analyses**

All behavioral, cognitive and immunohistochemical data were acquired and analyzed in a blind manner, in which the treatment conditions were blinded in the form of numerical codes. Likewise, all samples collected for LC-NSI-MS/MS analysis were randomly labeled by an experimenter before the measurements and analysis were conducted, with the samples being unblinded once all data were collected and analyzed. All statistical analyses were performed using SPSS Statistics (version 28.0, IBM, Armonk, NY, USA) and Prism (version 10.0; GraphPad Software, La Jolla, CA, USA). Statistical significance was set at $p < 0.05$. All data met the assumptions of normal distribution and equality of variance.

In the light-dark box test, the time spent in the light or dark compartment was analyzed by a 2 × 2 (treatment × compartment) repeated-measure ANOVA, whereas the total distance moved was evaluated using independent Student's *t*-test (two-tailed). In the open field test, the total distance moved, and the distance moved in the center zone were assessed using 2 × 5 (treatment × bins) repeated-measure ANOVAs. In the PPI test, % PPI and startle reactivity to pulse-alone trials were analyzed using 2 × 3 × 3 (treatment × prepulse intensity × pulse intensity) and 2 × 3 (treatment × pulse intensity) repeated-measure ANOVAs, respectively. All data in the Y-maze test were assessed using independent Student's *t*-tests (two-tailed). In the temporal order memory test, the absolute object exploration times and temporal order memory index were analyzed using 2 × 2 (treatment × object) repeated-measure ANOVA and independent Student's *t*-test (two-tailed), respectively. Group differences in the number of c-Fos positive neurons, % of c-Fos positive parvalbumin (PV) interneurons and total number of PV interneurons were assessed by independent Student's *t*-tests (two-tailed). Group differences in metabolites of the KYN pathway were assessed by independent Student's *t*-tests (two-tailed). All data involving PF-04859989 treatment were analyzed using one-way ANOVA. Whenever appropriate, ANOVAs were followed by Tukey's post-hoc test to control for multiple comparisons.

### SUPPLEMENTARY DATA

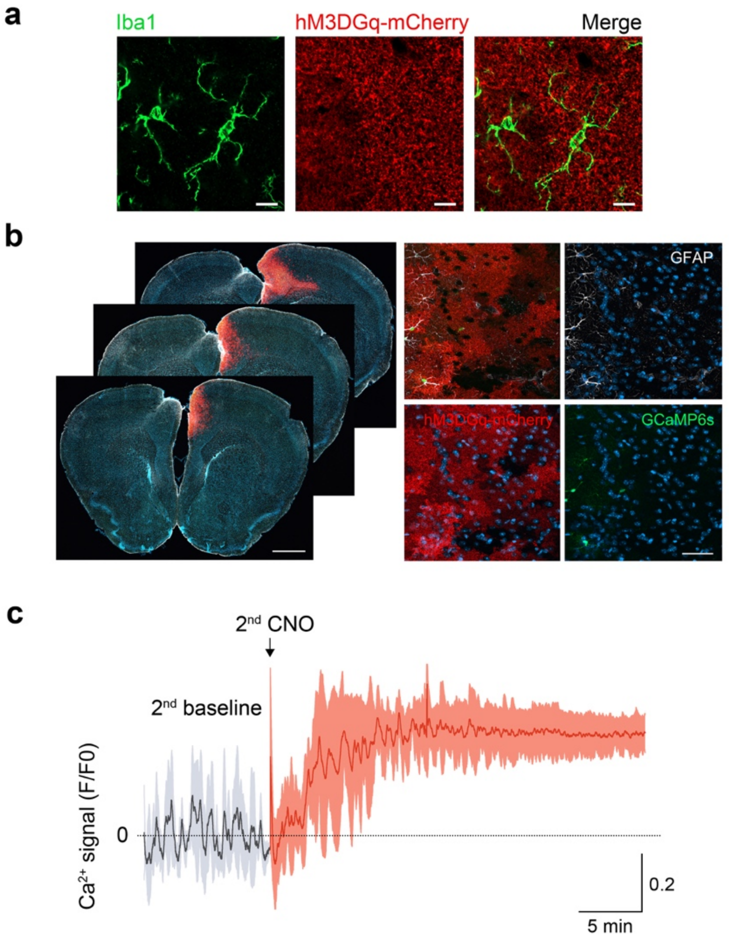


**Supplementary Figure S1.** Selectivity and effectiveness of hM3DGq-based stimulation of prefrontal astrocytes. **(a)** Representative double-immunofluorescence images confirming no hM3DGq expression (red) in microglia (Iba1, green). Scale bar = 10 µm. **(b)** Representative tile scan images of brain sections stained for mCherry (red), GFAP (white), and DAPI (blue) showing the location of microprism implantation (left). Photomicrographs on the right depict high resolution images showing the expression of hM3DGq-mCherry (red) and GCaMP6s (green) in prefrontal astrocytes in the hemisphere that has been imaged by two-photon microscopy. Note the absence of overt astrogliosis, as indicated by the low amount of GFAP expression (white). Scale bar = 1mm and 50 µm. **(c)** $Ca^{2+}$ response assessed by two-photon imaging in awake mice 48 hrs after the first imaging session and CNO challenge. Baseline $Ca^{2+}$ activity and $Ca^{2+}$ response to a second CNO treatment did not differ in prefrontal astrocytes (see main *Figure 1*). Black and red traces represent $Ca^{2+}$ responses (F/F0 ratio, means ± SD) during baseline and after 2$^{nd}$ CNO treatment, respectively. *N* = 4 male mice.

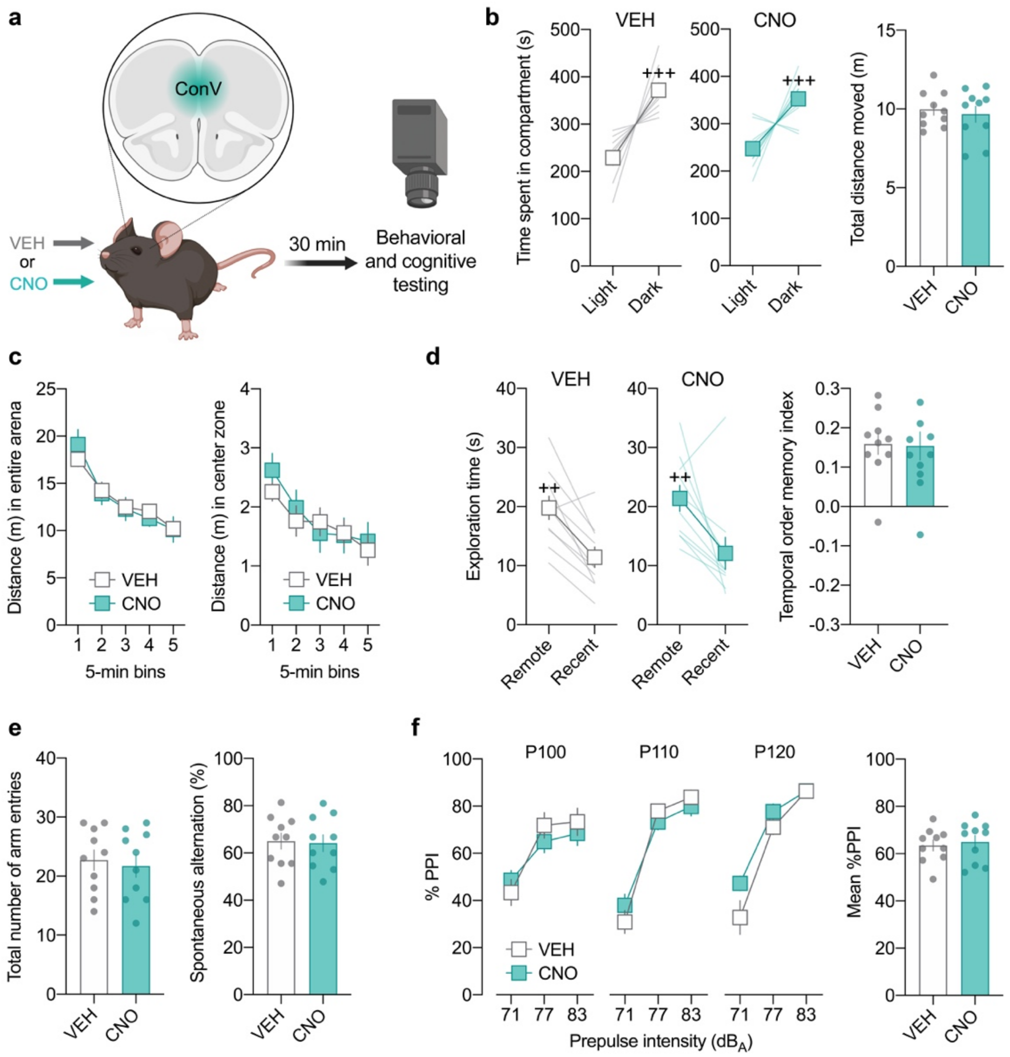


**Supplementary Figure S2.** Clozapine-N-oxide (CNO) treatment in ConV-expressing control mice does not alter behavior and cognition in control mice. **(a)** Male mice expressing a control AAV9 (ConV) in the prefrontal cortex were treated with vehicle (VEH) or CNO (1 mg/kg) and were then subjected to behavioral and cognitive testing 30 min after treatment. **(b)** Time spent in the light and dark compartments (line plots) and total distances moved (box plot) in the light-dark box test of innate anxiety-like behavior. $^{+++}p < 0.001$, reflecting the significant main effect of compartment revealed by repeated-measure ANOVA ($F_{(1,18)} = 37.9$). **(c)** Distance moved in the entire arena and center zone during the open field test of exploratory activity. **(d)** Absolute exploration times of the temporally remote and recent objects (line plots) and temporal order memory index (bar plot) in the temporal order memory test for objects. $^{++}p < 0.01$, reflecting the significant main effect of object in VEH-treated ($F_{(1,9)} = 10.3$) and CNO-treated ($F_{(1,9)} = 11.7$) mice revealed by repeated-measure ANOVA. **(e)** Total number of arm entries and percent spontaneous alternation in the Y-maze test of working memory. **(f)** Prepulse inhibition (PPI) test of pre-attentive filtering. The line plots show % PPI as a function of prepulse intensity (71, 77 and 83 $dB_A$) for each of the three pulse conditions (P100, P110 and P120, which correspond to pulse intensities of 100, 110 and 120 $dB_A$). The bar plot depicts the mean % PPI across all prepulse and pulse intensities. All data are means ± SEM with individual values overlaid; *N* = 10 mice per group and test.

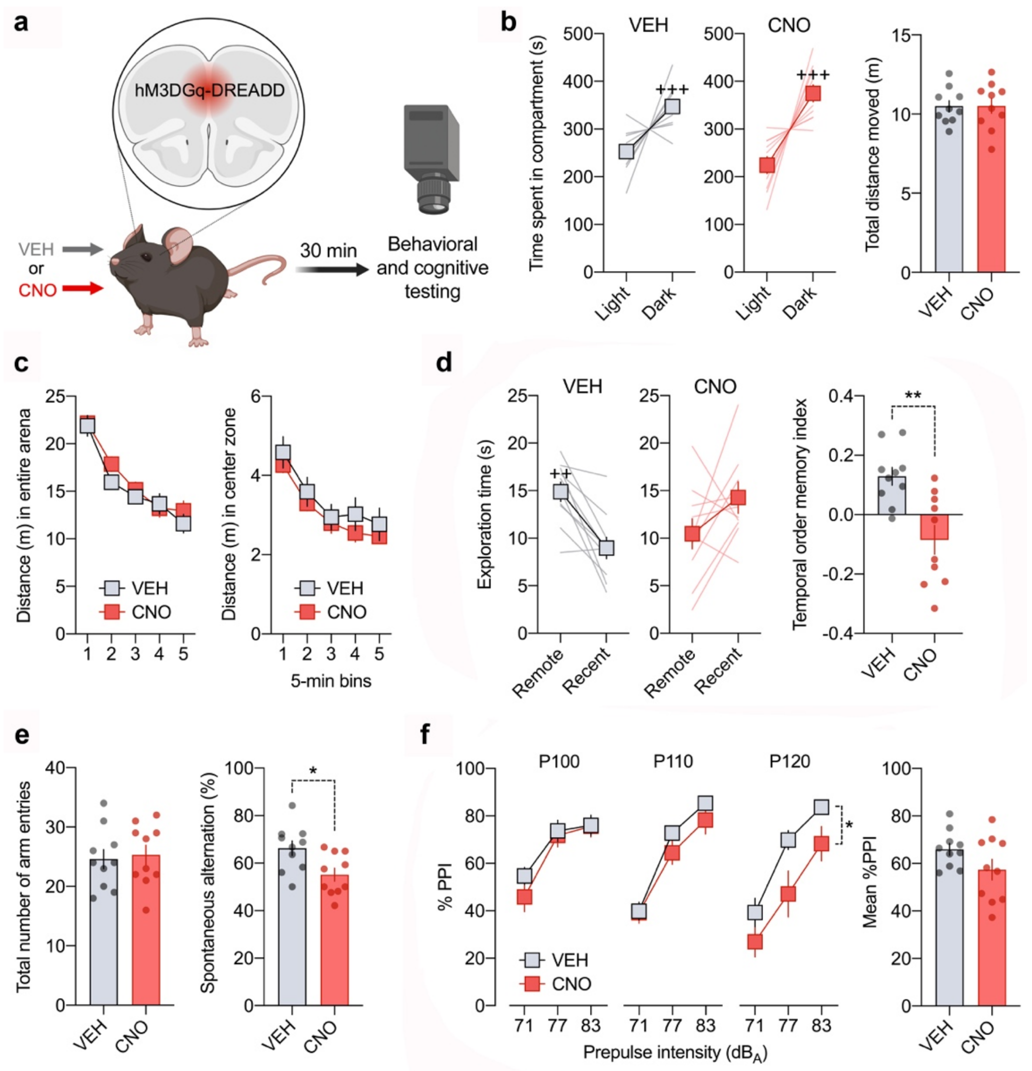


**Supplementary Figure S3.** Overactivation of prefrontal astrocytes impairs short-term memory and sensorimotor gating in female mice. **(a)** Female mice expressing hM3DGq in the prefrontal cortex were treated with vehicle (VEH) or clozapine-N-oxide (CNO, 1 mg/kg) and were subjected to behavioral and cognitive testing 30 min after treatment. **(b)** Time spent in the light and dark compartments (line plots) and total distances moved (box plot) in the light-dark box test of innate anxiety-like behavior. $^{+++}p < 0.001$, reflecting the significant main effect of compartment revealed by repeated-measure ANOVA ($F_{(1,18)} = 26.7$). **(c)** Distance moved in the entire arena and center zone during the open field test of exploratory activity. **(d)** Absolute exploration times of the temporally remote and recent objects (line plots) and temporal order memory index (bar plot) in the temporal order memory test for objects. $^{++}p < 0.01$, reflecting the significant main effect of object in VEH-treated mice revealed by repeated-measure ANOVA ($F_{(1,9)} = 15.4$); $^{**}p < 0.01$, based on two-tailed *t*-test ($t_{(18)} = 3.75$). **(e)** Total number of arm entries and percent spontaneous alternation in the Y-maze test of working memory. $^{*}p < 0.05$, based on two-tailed t-test ($t_{(18)} = 2.70$). **(f)** Prepulse inhibition (PPI) test of sensorimotor gating. The line plots show % PPI as a function of prepulse intensity (71, 77 and 83 dBA) for each of the three pulse conditions (P100, P110 and P120, which correspond to pulse intensities of 100, 110 and 120 dBA). The bar plot depicts the mean % PPI across all prepulse and pulse intensities. $^{*}p < 0.05$, reflecting the significant main effect of treatment in the P120 condition, as revealed by repeated-measure ANOVA ($F_{(1,18)} = 5.19$) in the P120 condition. All data are means ± SEM with individual values overlaid; *N* = 10 female mice per group and test.

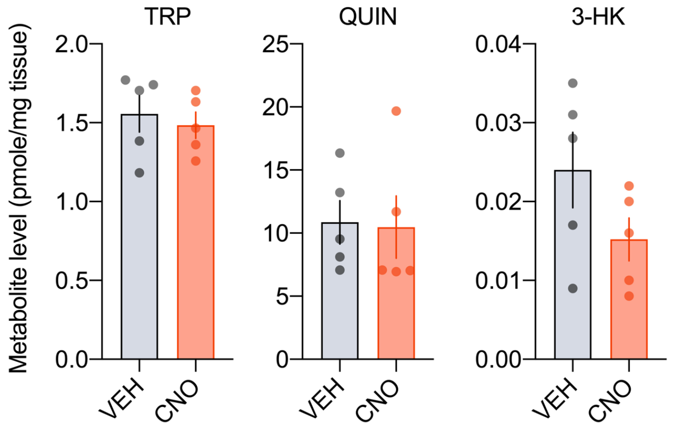


**Supplementary Figure S4.** Levels of tryptophan (TRP), quinolinic acid (QUIN), and 3 hydroxykynurenine (3-HK) in the prefrontal cortex of hM3DGq mice after vehicle (VEH) or clozapine-N-oxide (CNO, 1 mg/kg) treatment. Each data point represents the pooled samples of two mice, with 5 replications for each group and measurement. All data are means ± SEM, with individual data points overlaid.

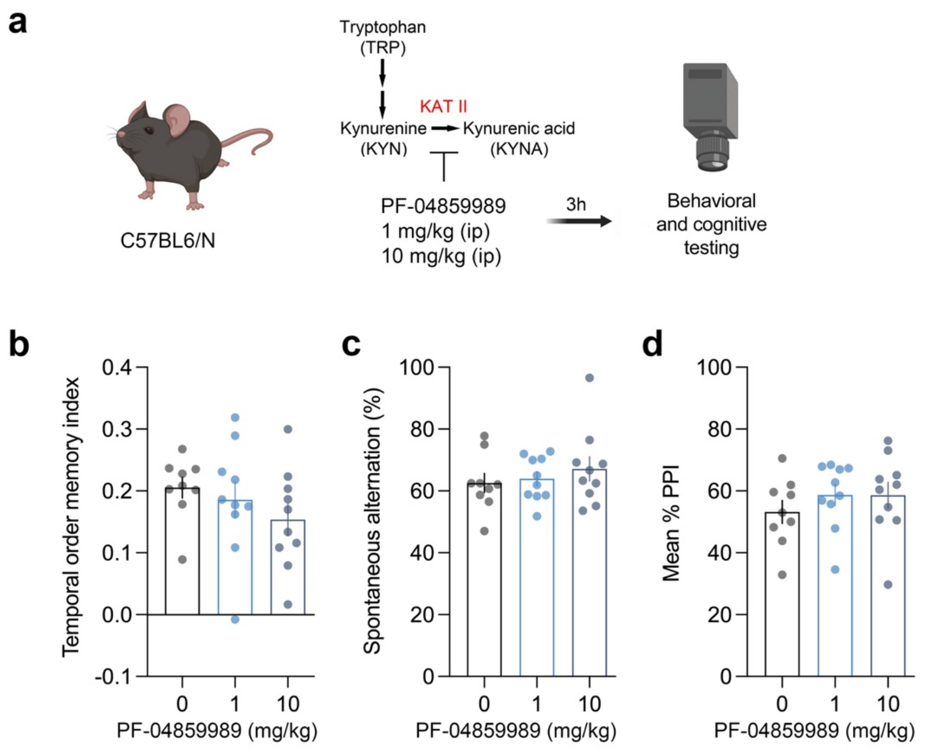


**Supplementary Figure S5.** Pharmacological inhibition of central KYNA in C57BL6/N male mice does not alter short-term memory and pre-attentive filtering under basal (i.e., non-DREADD) conditions. **(a)** Male C57BL6/N mice received 0, 1 or 10 mg/kg PF-04859989 and were subjected to behavioral and cognitive testing 3 hs after treatment. **(b)** Temporal order memory index in the temporal order memory test for objects. **(c)** Percent spontaneous alternation in the Y-maze test of working memory. **(d)** Prepulse inhibition (PPI) test of pre-attentive filtering. All data are means ± SEM, with individual data points overlaid. *N* = 9 (0 mg/kg) and *N* = 10 mice per remaining groups.
